# Responses of root-associated fungal communities of mature beech and spruce during five years of experimental drought

**DOI:** 10.1101/2023.09.24.559161

**Authors:** Fabian C. Weikl, Thorsten E. E. Grams, Karin Pritsch

**Author notes:** Author for correspondence: Fabian Weikl.

## Abstract

Drought affects the fine-root systems of European beech (*Fagus sylvatica* L.) and Norway spruce (*Picea abies* [L.] KARST) in different ways, but little is known about how this impacts their fine-root-associated fungal communities.

In a five-year throughfall exclusion experiment (KROOF) in a mature stand, we investigated whether recurrent drought periods progressively alter fine-root associated fungal communities, fine-root vitality, and ectomycorrhizal functionality in relation to the tree root zone (pure beech, pure spruce, or their mixture) and abiotic soil parameters.

We found that the influence of recurrent droughts on root fungal communities peaked in the third year of the experiment and affected fungal functional groups in different ways. The root zone was the predominant factor in structuring all functional groups of root-associated fungi, while we did not find a prominent effect of root mixture. The importance of other factors (year of sampling, soil depth) varied among fungal functional groups.

Our results indicate a robust biotrophic root-fungal system relying mainly on surviving root tips, complemented by a fluctuating saprotrophic fungal assembly.

## Introduction

Drought and heat waves have increased forest tree mortality worldwide (Allen et al., 2010; Hartmann et al., 2022). In Central Europe, tree species such as Norway spruce, which has been planted outside its natural range on a large scale, and European beech, among others, suffered particularly from extreme drought events (Caudullo et al., 2019; Schuldt et al., 2020; Obladen et al., 2021).

However, long-term observations of stand growth revealed a positive effect in mixtures of beech and spruce with an average of 8 % higher yield compared to monospecific stands (cf. Pretzsch et al., 2010, 2013, 2020). Synergies of both tree species are said to be different rooting depths (shallow-rooted spruce vs deeper-rooted beech), asynchrony of water use and photosynthesis by evergreen vs deciduous foliage (cf. Pretzsch et al., 2014; Goisser et al., 2016), differences in litter turnover (Albers et al. 2004), and soil nutrient cycling (Achilles et al. 2021).

Both species react differently to drought. Spruce, as isohydric species, minimises water loss under drought at the expense of photosynthetic activity (Hartmann et al., 2013), while beech, as a more anisohydric species, with lower stomatal control under drought, is more prone to water loss while maintaining photosynthetic activity (Leuschner, 2020). Despite their frequently observed vulnerability to drought, results from the Kranzberg roof (KROOF) experiment showed that both tree species can successfully adapt to extreme drought, at least temporarily (Grams et al., 2021). In this experiment, exclusion of precipitation throughfall for five consecutive vegetation periods resulted in tree death only in few cases, although soil desiccation was progressive and reached deeper soil layers already in the second year (Pretzsch et al., 2020; Grams et al., 2021). Both tree species responded to the extreme drought with greatly reduced above- and belowground growth (Grams et al., 2021; Jacobs et al., 2021).

Soil desiccation changes multiple processes in the overall plant-soil system. Ultimately, however, it is the lack of connection to soil water that leads to tree desiccation. The nutrient- and water-absorbing fine roots of trees provide the connection between soil water and xylem and are, therefore, crucial for the water supply of trees (Körner et al., 2019). Under drought, spruce maintains absorbing fine roots during dry periods and protects them from desiccation by suberisation, while beech continuously forms new fine roots with a comparatively short life span (Nikolova et al., 2020). Fine roots of both tree species can associate with numerous species of ectomycorrhizal fungi (EMf) – at the KROOF site, 144 phylotypes were detected previously (Nickel et al., 2018). Through their extramatrical hyphae, EMf connect to soil where they forage for nutrients (Read et al., 2003; Pritsch & Garbaye 2011; Lindahl & Tunlid 2015) and connect to soil water (Lehto & Zwiazek 2011). Spruce EMf-community composition has been shown to depend on neighbouring tree species (Otsing et al., 2021). In view of recent findings that tree growth, and EMf compositions are correlated (Anthony et al., 2022), tree mixture effects of spruce, for example with beech, could also play a role in tree performance under drought (Pretzsch et al., 2020).

In addition to EMf, which dominate root-associated fungi in terms of biomass, other fungi are associated with ectomycorrhizae and their mycorrhizosphere, such as saprotrophic or pathotrophic fungi and many whose functions are still unknown (Tedersoo et al., 2009; Mestre et al. 2021; Wang et al., 2020).

Soil drought affects soil microorganisms in complex ways (Schimel et al., 2018), leading to a slowdown in biological processes as drying progresses and eventually to a halt in extreme drought (Manzoni et al., 2012). Accordingly, the enzymatic degradation of organic compounds is strongly reduced, leading to an accumulation of litter (Bastida et al., 2017). Simultaneously, the uptake of nutrients by soil microorganisms and plants is reduced, leading to, e.g., the accumulation of nitrogen compounds, particularly NH ^+^, due to limited diffusion and hampered nitrification (Stark & Firestone 1995). This, in turn, can influence soil fungal community composition (Cline et al. 2018) and is additionally modulated by the position in the soil profile (Weber et al. 2013).

In a study on the first three years of the KROOF experiment, we showed a significant and increasing change of the α-diversity and composition of EMf communities on fine roots of spruce and beech from throughfall-exclusion (TE) plots (Nickel et al., 2018). This was interpreted as an adaptation to increasing soil dryness and loss of fine roots over the years. At the same time, however, there was no significant change in the activities of hydrolytic enzymes of EMf in TE plots compared to controls (CO) (Nickel et al., 2018), which can be interpreted as high functional redundancy and thus resistance at the level of the overall EMf community (Courty et al., 2005). Nevertheless, the severe loss of absorbing fine roots resulted in a strong decrease of vital mycorrhizal tips and, consequently, a significant reduction of potential enzyme activities at the ecosystem level (Nickel et al., 2018).

In the present study, we examined the development of the entire root associated fungal communities (symbiotrophic, saprotrophic, pathotrophic) over five consecutive years of the KROOF experiment. For a complete picture, we included the dataset on EMf from the first three years (published in Nickel et al. 2018). We included soil moisture and nitrogen contents as environmental parameters, and enzyme activities on vital EM were determined as a measure of EM function. Our data analysis relates to the following hypotheses:

(H1) TE over multiple growing seasons will progressively alter the diversity, composition and functional traits of root-associated fungi of TE trees compared to CO trees, continuing trends for EMf from the three-year drought.

(H2) Reduced numbers of vital roots under TE will be associated with a progressively lowered α-diversity of biotrophic fungi (EMf, endophytes, pathogens) over time and result in changed communities. This effect is more pronounced in spruce than in beech because of their different fine root growth under drought (2a).

(H3) Low soil moisture will reduce α-diversity and consequently strongly change saprotrophic fungal communities.

(H4) Niche diversification through tree mixing creates an advantage over monospecific root zones under TE and leads to higher fungal diversity and more shared taxa (H4a), including more efficient soil exploration (exploration types of EMf) (H4b).

## Materials and Methods

### Site description and throughfall-exclusion

The field experiment KROOF has been described in detail by Pretzsch et al. (2014) and Grams et al. (2021). In brief, it is a spruce-beech forest in Southern Germany (11°39’42’’E, 48°25’12’’N; 490 m a.s.l.), planted in 1951 ± 2 AD (Norway spruce (*Picea abies* [L.] KARST)) and 1931 ± 4 AD (European beech (*Fagus sylvatica* L.)), respectively in a nutrient-rich luvisol. 12 plots between 100-200 m² were established in 2010 by trenching to 1 m depth. A water-impermeable tarp was inserted around the plots down to the trenching depth to minimise lateral water permeation. Plots were fitted into the existing forest in a way that each plot contained a group of 3-7 mature specimens of both tree species, resulting in root zones with spruce trees neighbouring spruce (ss – pure spruce root zone), beech trees neighbouring beech (bb), and an interspecific contact zone (mixture – m) containing beech (bm) and spruce roots (sm). In 2013, retractable roofs were installed below the tree canopy on six out of 12 plots to allow precipitation throughfall-exclusion (TE) by automatically closing during periods of rainfall. Roofs remained open during winter to allow for refilling of the soil water reservoir. The other six plots served as controls (CO). Plots were designed pairwise, with one CO next to one TE plot being an experimental unit. Air temperature and precipitation were permanently recorded; volumetric soil water content was assessed weekly on each plot and in each root zone at four different depths using time domain reflectometer (TDR) probes.

As detailed in Grams et al. (2021), TE resulted in an almost complete non-availability of soil water in the upper 70 cm during the summer periods of 2014-2018, with volumetric soil water contents of TE being consistently about 10% lower than in controls in all soil layers. While soil water content in TE plots was close to the permanent wilting point of the trees in all summers except 2014, such a situation was only found during the summer of 2015 for CO, as a result of an exceptional natural summer drought coupled with low annual precipitation and high vapour pressure deficit (Grams et al., 2021). Soil water contents (0-7 cm) and the periods of activated roofs are given in Fig. 1a.

**Figure 1:**
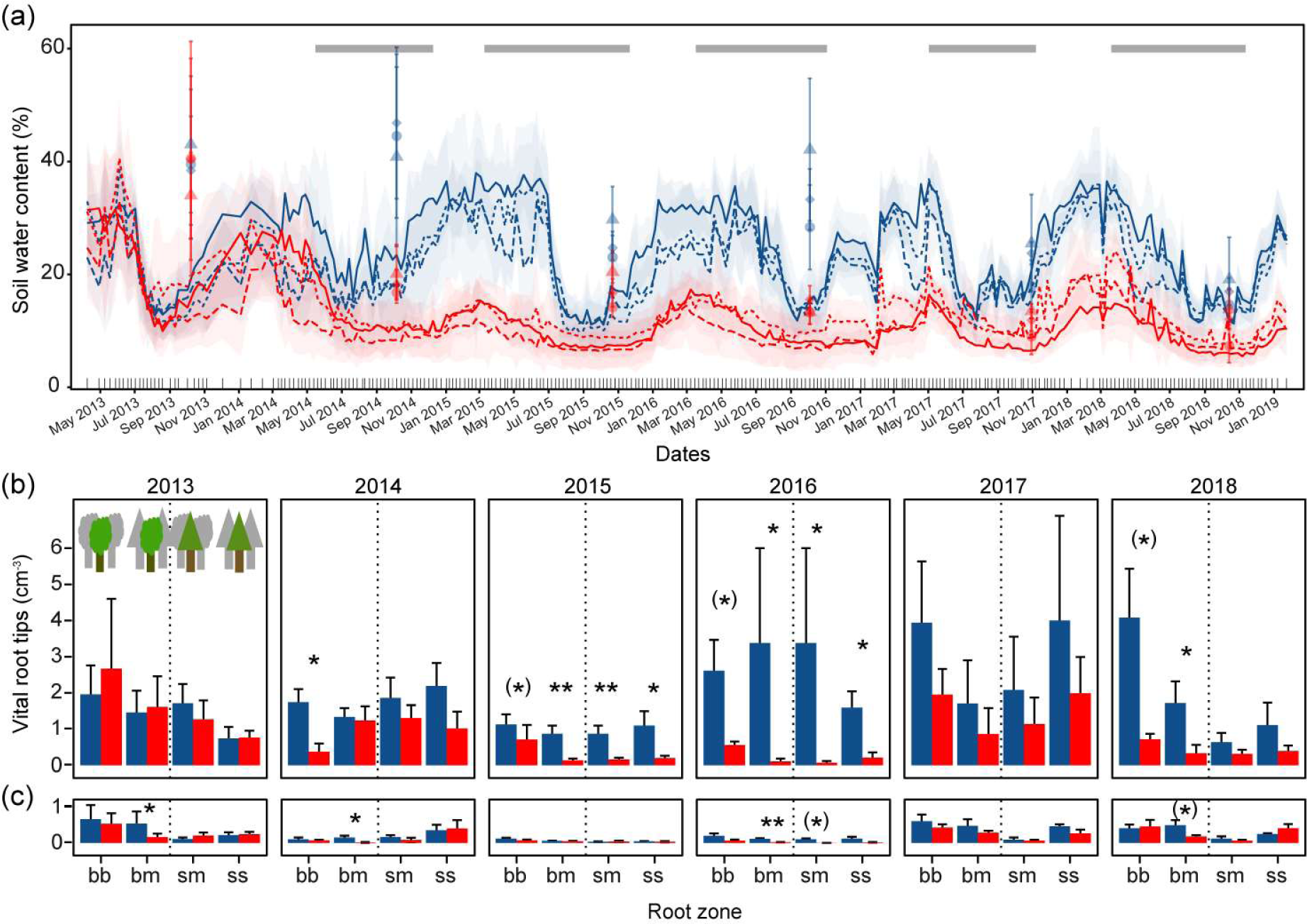
Soil water contents (SWC) and vital root tips before (2013) and during the throughfall-exclusion experiment (TE). Red: TE, dark blue: controls (CO). a) SWC in the upper soil layer (c. 0 - 7 cm). Lines with sd-ribbons: volumetric SWC measured with stationary time domain reflectometers (TDR) (full: beech, dashed: spruce, dotted: mixture); symbols: gravimetric SWC of the sampled soil originating from different tree root zones (triangles: spruce, diamonds: mixture, circles: beech) with standard deviations (sd); data partly reported earlier (Nickel et al. 2018, Grams et al. 2021); marks along the x-axis: TDR measurement dates; grey horizontal bars: TE periods. b, c) numbers of vital root tips of per soil volume in the upper (c. 0 - 7 cm) (b) and lower soil layers (c. 7 – 30 cm) (c); bb: pure beech root zone, bm: beech roots in tree mixture root zone, sm: spruce roots in mixture root zone, ss: pure spruce root zone, error bars: ± 1 standard error; Wilcoxon p: (*): < 0.1, *: < 0.05, **: < 0.01, nunits = 4-6 (two experimental units removed after 2015 due to bark beetle attack)

### Root, mycorrhiza, and soil sampling

Sampling was performed annually in late autumn before opening the roofs for the winter period. It started in 2013 (before TE) and continued until 2018. Each time, soil cores of 4 cm diameter were taken to a depth of 25–40 cm, divided into an upper organic (horizons Of + hAh) and lower mineral layer (AlBv), their respective thickness recorded, and collected separately. Samples were kept at 4-6 °C for up to 4 weeks until further processing. On every plot, one soil core was taken from zones ss and bb, and two from m.

### Soil parameters

Root-free subsamples were taken in triplicates before root sorting and used for gravimetric soil water (5 g fresh soil) and nitrogen content (3 g fresh soil). Soluble inorganic nitrogen (N_min_) was determined using extracts (0.01 M CaCl_2_/soil = 1/4 w/v, 45 min overhead shaking, filtration through Whatman filter paper 595½) and an automatic analyser with continuous flow air segmented injection (SA20/40, Type 5100; Skalar Analytical, Breda, Netherlands) (VDLUFA 1991).

### Fine root parameters

Roots were sorted from soil samples with forceps, placed in tap water and stored overnight at 4°C. The next day and within 30-60 min per sample, fine roots (diameter <1 mm for beech, <2 mm for spruce) were assigned to beech and spruce, all vital tips were counted and assigned to EM morphotypes under a stereomicroscope. From each root sample, 21 vital EM representing the most abundant morphotypes (n ≥ 3) were used for extracellular enzyme activity measurements (Supporting Information (SI) Method S1). Cleaned fine roots were gently dried with a paper tissue and frozen at -20 °C.

### Sample processing for amplicon sequencing

480 fine root samples [6 years × 2 soil layers × 12 (6 CO, 6 TE) plots × 4 root zones (pure zones (bb, ss), beech (bm) and spruce (sm) roots from the mixture zone] were processed (two experimental units removed after 2015 due to bark beetle attack). DNA extraction followed Nickel et al. (2018): In brief, DNA from c. 100–400 mg of fine root homogenate was extracted with PowerSoil DNA Isolation Kits (Qiagen, Hilden, Germany) with modifications as described therein.

ITS2 rDNA was amplified with PCR primer mixes for maximum phylogenetic recovery (Tedersoo et al., 2015; Table S1) with adapters for the Miseq workflow (protocol #15044223 Illumina, San Diego, CA, USA). Reactions contained 1 µl DNA (5 ng, except for extraction-negative-controls which contained 1 µl undiluted extract, and PCR-negative controls), 0.5 µl ITS3mix (10 pmol equimolar mix of ITS3-Mix1 to -Mix5), 0.5 µl ITS4mix (10 pmol equimolar mix of ITS4-Mix1 to -Mix4), 10 µl NEBNext High-Fidelity 2X PCR Master-Mix (New England Biolabs, Frankfurt, Germany), 8 µl H_2_O. PCR profiles: 5 min 95° C, 28 × [30 s 95 °C, 30 s 55 °C, 60 s 72 °C], 10 min 72 °C. The quality of all products was checked on agarose gels.

Triplicate PCRs were pooled and cleaned using Agencourt AMPure XP (Beckman Coulter, Krefeld, Germany) with a bead:DNA ratio of 1:1. Removal of primers was controlled with a Fragmentanalyzer DNF-473 kit (Agilent Technologies, Waldbronn, Germany) and yield quantified using fluorescent-dye-based assays.

Eight PCR cycles with individual dual-index combinations of Nextera XT Index v2 kits (Illumina) were performed (1 µl DNA (5 ng), 2.5 µl index 1 (i7 series), 2.5 µl index 2 (i5 series), 12.5 µl NEBNext High-Fidelity 2X PCR Master Mix, 6.5 µl H_2_O). Indexed amplicons were cleaned, size-checked and quantified as above. Amplicons (4 nM) of each sample were pooled. Final preparations and sequencing (Miseq v3, 600 cycles flowcells, Illumina) followed the manufacturer’s recommendations (protocol #15044223 Rev. B).

### Data origins

Lab work for the first part of the time course (until 2016) was done within the preceding study (Nickel et al., 2018). Root vitality and EMf-related data on this part are shown therein and used here to cover the complete drought period.

### Processing of sequences

Raw data from the previous study and newly generated data were processed together using PIPITS fungal ITS analyses pipeline v2.7 (Gweon et al., 2015) following the reference workflow: joining read-pairs with VSEARCH v2.15.0 (Rognes et al., 2016) (parameters: -q 30; Q 33, p-value of assembly ≤ 0.0001), FASTQ_QUALITY_FILTER (parameters: -q 30, -p 80; Q33; FASTX-Toolkit v0.0.14, https://github.com/agordon/fastx_toolkit)), extraction of ITS2 with ITSX v1.1b (Bengtsson-Palme et al., 2013), removal of sequences < 100 bp and clustering of OTUs by 97% sequence identity (VSEARCH), chimera removal using UNITE UCHIME reference data (v7.2, Nilsson et al., 2015), mapping of reads onto OTUs, removal of singletons, assignment of taxonomy with RDP Classifier (rdptools v2.0.2) (Wang et al., 2007) against UNITE 8.2 (Abarenkov et al., 2020). All analyses of HTS data were done on phylotype tables, which resembled data structure obtained from EM enzyme activities and mophotyping better than OTUs (Nickel et al., 2018). Taxonomic assignments with low confidence (<0.85) were omitted.

Information on primary fungal lifestyles (‘lifestyle’ in the following) was obtained from FungalTraits (Põlme et al., 2021). EMf and exploration types were manually reviewed, guided by Agerer (2001) and Tedersoo & Smith (2013), which only resulted in minor changes (new ‘EMf-borderline’ sub-group).

### Statistical analyses

Values of p < 0.05 were treated as significant. Non-parametric tests were used when normality or homoscedasticity of variances was not met. Analyses were conducted in R v4.0.3 (R Core Team, 2020).

Samples with low sequencing depth (< 9500 sequences) and non-fungal phylotypes were removed. Reads were 10^4^ times randomly rarefied using GUniFrac (Chen, 2012), and the results averaged to compare all samples at equivalent sequencing depths (Weiss et al., 2016; Cameroon et al., 2021). Bray—Curtis dissimilarities between samples and biodiversity indices were calculated using vegan (Oksanen et al., 2017), figures produced with base R, vegan, ggplot2 (Wickham, 2016), ampvis2 (Andersen et al., 2018), and phyloseq (McMurdie & Holmes, 2013). Effects of environmental parameters on communities were tested using Permutational Multivariate ANOVA Using Distance Matrices (PERMANOVA) on Bray—Curtis dissimilarity matrices (vegan 2.5-7 adonis(); 10^5^ permutations).

## Results

### Soil parameters and vital root tips

Volumetric soil water contents in the upper 7 cm on TE plots were always below levels of CO during the drought phases (Fig. 1a, details: Grams et al., 2021). Ammonium and N_min_ gradually rose in CO and TE upper soil (but peaked in 2016 for TE spruce), while nitrate rose in CO but not continuously in TE (Fig. S1). The majority of vital root tips of both tree species occurred in the upper soil (Fig. 1b) and did not shift towards the lower soil for TE compared to CO (Fig. 1c). Numbers of vital root tips per soil volume in each tree rooting zone started at similar levels in plots assigned to subsequent CO and TE treatments (2013) and decreased in TE relative to CO plots during the years of treatment (2014-18). Differences between CO and TE were significant in 2015 and 2016 for most root zones (Fig. 1b) but rarely later (2017-2018). Numbers of vital root tips varied between years (Figs. 1a,b). Influences of tree mixture were not resolved (high variation between samples, especially in 2016 and 2017; Fig. 1b).

### Root fungal diversity

ITS2 sequencing of 468 samples yielded 21 × 10^6^ quality-filtered fungal reads without singletons (average sample^-1^: 45195, median: 40849) assigned to 4853 OTUs and 1625 phylotypes (1060 phylotypes > 10 sequences).

Diversity indices of the total root-associated fungal community (Fig. S2) and the major (see below) fungal lifestyle categories EMf, saprotrophs, and unknown lifestyle did not differ between CO and TE or throughout the time course (Figs. 2, S2). An exception was the pure spruce root zone, where EMf diversity from the upper soil was significantly lowered under TE (Shannon 2015-2017, Simpson 2015-2017, Evenness 2016).

**Figure 2:**
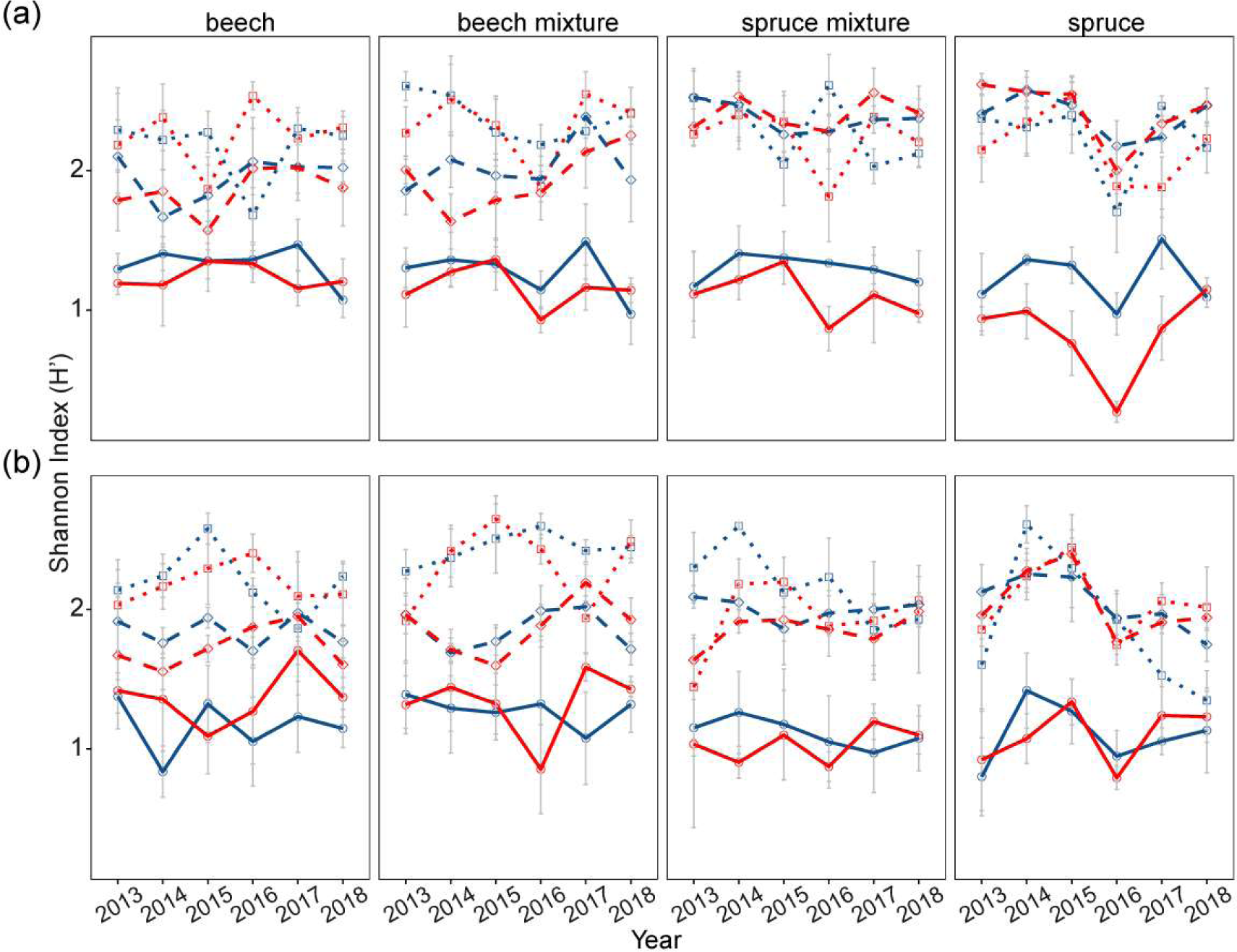
Shannon diversity of the most important fungal lifestyles in the data-set divided by their growth in pure or mixture root zones of the tree species. Solid line: EMf, dashed: unknown lifestyle, dotted: saprotrophs; blue: control (CO) plots, red: throughfall-exclusion (TE), error bars: ± 1 standard error. Only samples from plots were used that persisted during the whole experiment (nunits = 4). 2013 shows the values before the start of TE. a) upper soil, b) lower soil. Significant differences CO/TE in upper soil (Wilcoxon p < 0.05): 2016 pure beech saprotrophs, 2016 mixture spruce EMf, 2015-2016 pure spruce EMf, 2017 pure spruce saprotrophs; in lower soil: 2013 unknown mixture spruce, 2014 saprotrophs mixture spruce,

### Annually accounted TE effects on major fungal lifestyles

EMf communities in TE increasingly diverged from CO until the third year of experimental drought (2016), when the effect size (PERMANOVA R^2^_adj_) of the factor *treatment* was higher than the influence of the factors *root zone* and *soil layer* (Fig. 3). This trend reversed for the last two years.

**Fig. 3:**
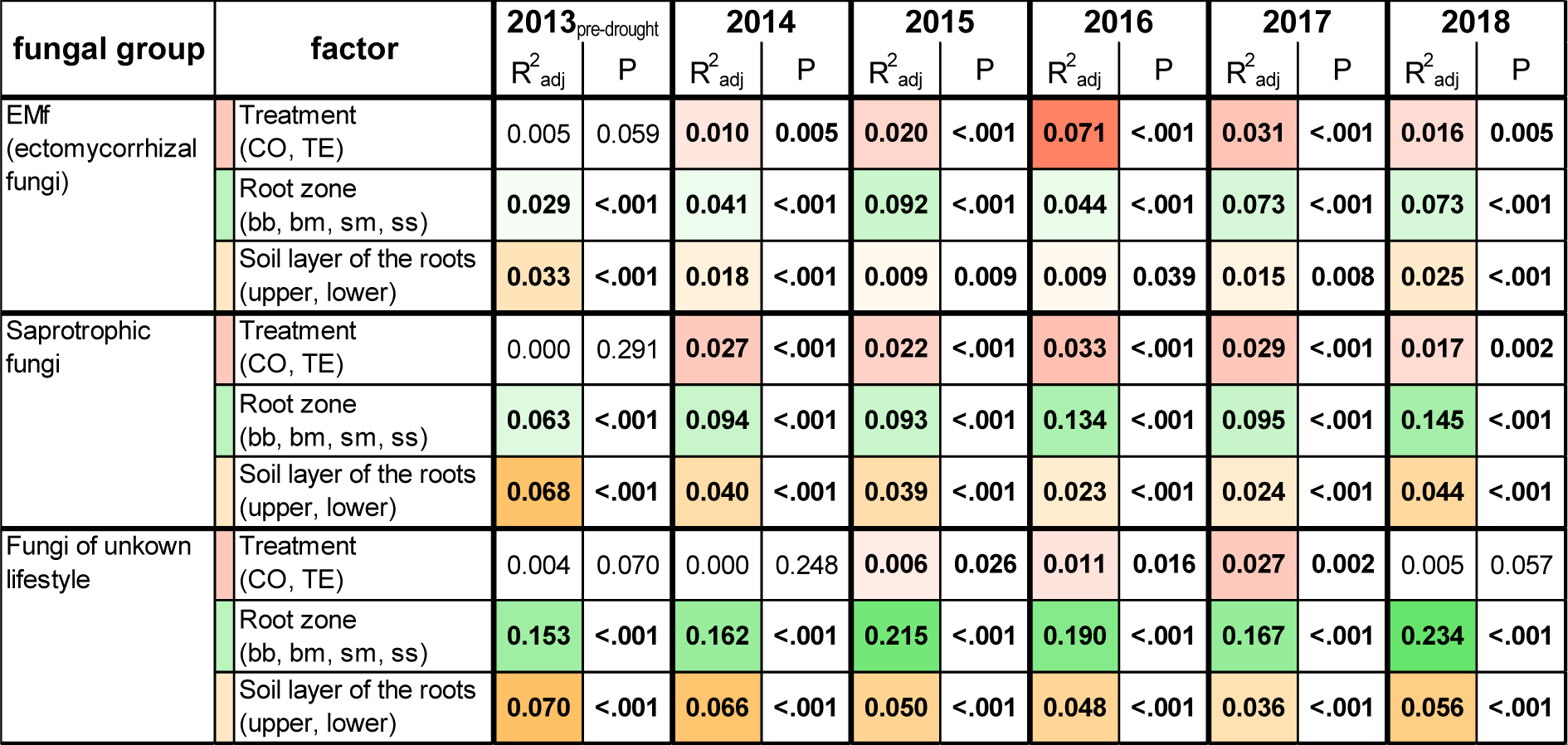
Annual PERMANOVA results on major experimental factors for important fungal groups. *Treatment* CO: controls, TE: throughfall-exclusion. *Root zone* bb/ss: roots from monospecific zone of beech/spruce, bm/sm: beech/spruce roots from root mixture zone. *Soil layer of the roots*: upper: c. 0-7 cm, lower: c. 7-25 cm. Bold: significant (P < 0.05) results. Colour intensities: highest to lowest significant result per tested effect. P: calculated from Bray-Curtis dissimilarities using adonis() with 10^4^ permutations and *experimental unit* as random and blocking factor; R^2^adj: adjusted R^2^ calculated using vegan::varpart(). 2013-2015: nunits = 6, 2016-2018: nunits = 4). Significant interactions were found for saprotrophs (2013-2014: *root zone* × *soil layer*; 2016: *treatment* × *root zone*) and unknown lifestyles (2014: *treatment* × *soil layer*; 2015: *treatment* × *root zone*, *root zone* × *soil layer*; 2016: *root zone* × *soil layer*). A comparison with nunits=4 for the whole time series is given in Table S2.

For community composition of saprotrophic fungi, the effect of *treatment* was significant from the first year, too. However, effect sizes stayed similar between years, remained smaller than for *root zone*, and exceeded that of *soil layer* just in 2016 and 2017 (Fig. 3).

For fungi of unknown lifestyle, the effect of *treatment* was only significant in 2015-2017, during which it increased (Fig. 3). Noteworthy, the influence of *root zone* was very pronounced (R^2^ 0.153-0.234) during the whole observation period.

### Overall effects of five years TE on fungal communities with different lifestyles

In a combined analysis of CO and TE communities from all treatment years, the three major groups EMf, saprotrophs, and fungi of unknown lifestyles showed three distinct reaction patterns (Fig. 4). EMf showed a clear effect of TE in unconstrained ordination (Fig. 4a) without progression of community dissimilarities throughout the years for CO and TE and samples from pure beech and spruce zones being most dissimilar. Therein, significant correlations with environmental vectors were found with N_min_ towards spruce, soil water contents at the time of sampling, and vital root tips cm^-^³, the latter two towards CO. This is reflected in the PERMANOVA model for EMf (Fig 4b), with the biggest effect for the factor *root zone*, followed by *treatment*, *soil layer,* and only a moderate effect for *year of treatment*. The relation to *N_min_* was significant with a small effect size, while the relation to *vital root tips cm*^-^*³ soil* remained insignificant (Fig. 4b).

**Figure 4:**
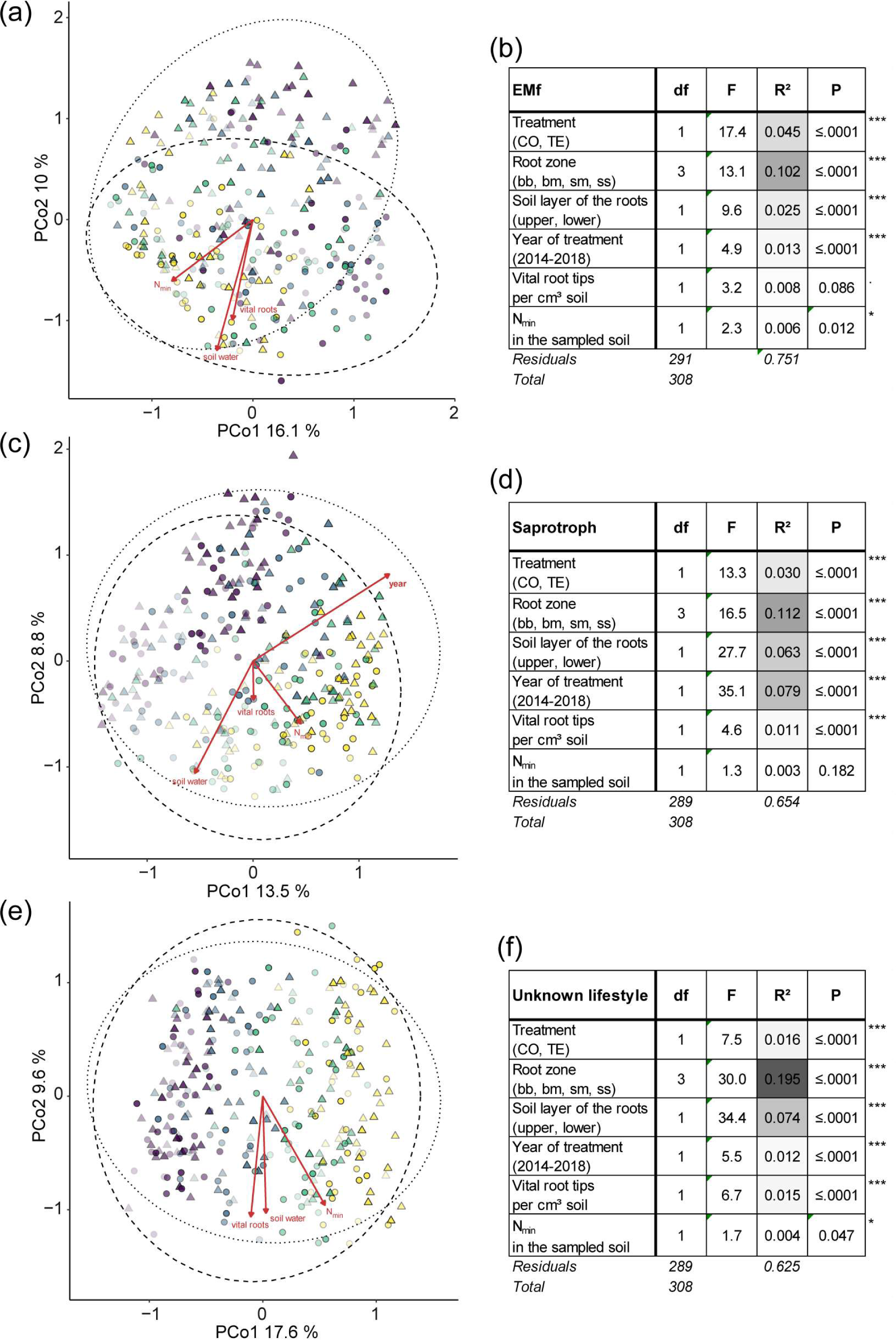
Variation of root fungal communities and influence factors during five years repeated drought as seen through principal coordinate analysis (a, c, e) and PERMANOVA (b, d, f) for EMf (a, b), saprotrophs (c, d), and fungi of unknown lifestyle (e, f).

Saprotroph communities did not show a clear influence of TE in the ordination (Fig. 4c). However, CO and TE communities changed over the years, highlighted by a significant correlation vector for the sampling year and an almost opposing significant vector for soil water content. N_min_ pointed towards samples from spruce. PERMANOVA indicated a *root zone* effect of similar size as seen for EMf, a prominent influence of *soil layer*, and a smaller *treatment* effect (Fig. 4d). The effect of *year of treatment* was much stronger for saprotroph than for EMf communities (R² 0.079 vs 0.013). In addition, a minor significant effect was found for *vital root tips cm*^-^*³* but not for *N_min_* (Fig. 4d).

Fungi of unknown lifestyle gave a distinctive picture in the ordination (Fig. 4e), without evident fractionation by TE or the year of treatment, but a precise separation by the root zone along the first axis, including a separation of pure vs both mixture zones. Significant correlation vectors for soil water at the time of sampling and vital root tips cm^-^³ were aligned in parallel to the second ordination axis. N_min_ was significantly correlated with spruce. Reflecting the ordination, PERMANOVA indicated a very pronounced relationship with the factors *root zone* (R = 0.195) and a still important effect of *soil layer* (R² = 0.074), while *treatment*, *year of treatment* and *vital root tips cm*^-^*³* only had moderate-minor effects (Fig. 4f).

Samples are depicted by triangles (TE – throughfall-exclusion) and circles (CO – controls). Colours display the *root zone*: lilac: beech roots in beech monospecific zone (bb), blue: beech roots in tree mixture zone (bm), green: spruce roots in tree mixture zone (sm), and yellow: spruce roots in spruce monospecific zone (ss). Colour-intensity varies according to sampling years: darkest 2018 (5th year TE), lightest 2014 (1st year TE). 95% confidence ellipses are given for TE (dotted lines) and CO (dashed). Arrows: significantly fitted environmental vectors (year = year of TE, Nmin = mineral Nitrogen content in the corresponding soil, vital roots = vital root tips cm^-^³ soil, soil water = gravimetric soil water content in the corresponding soil samples; correlation to Nmin is non-linear, other correlations linear or monotonous). Ordinations and PERMANOVAs were calculated from Hellinger-transformed data using Bray–Curtis dissimilarities and only samples from those plots were used which persisted the whole observation period (nsamples = 310, nexperimental units= 4). PERMANOVAs (one sample with incomplete data excluded) were done with 10^4^ permutations using experimental units as random and blocking factor. Full models information including significant interactions are given in Table S3.

### Major root-associated orders

The top 10 fungal orders (Fig. 5a) comprised 71-85 % of the relative shares, depending on root zone and treatment. Helotiales dominated (> 22%) without a clear preference for a soil layer or root zone or response to TE (Fig. 5a). Russulales were especially abundant in pure beech (up to 27 %) and much less in pure spruce zones (<10%). Similar tendencies were found for Agaricales (11-20% vs 4-13%) and Mortierellales (4.7-7.8% vs 1.8-6.2%). Boletales were most abundant in beech mixture. Archaeorhizomycetales had a clear preference for spruce roots in the lower soil of CO (10.4%) and were diminished there with TE (5.4%) (Fig. 5a). Similarly, Atheliales (2.7-4.7% in) were reduced in TE (0.3-2%).

**Figure 5:**
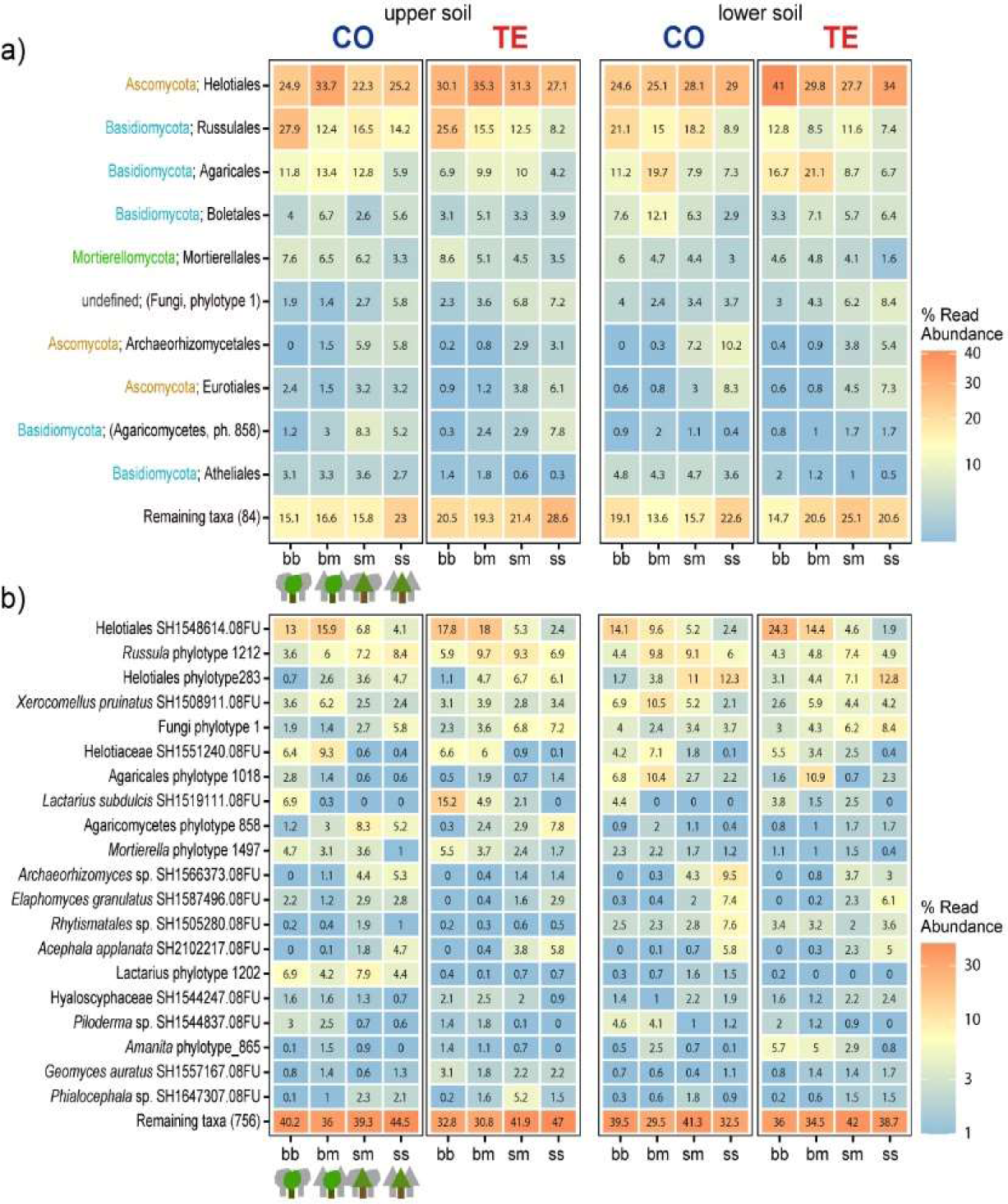
Most abundant root-associated fungal taxa and their shares under different conditions averaged for the treatment years (2014-2018). a) top 10 fungal orders (average of relative abundance per sample) under different conditions, n = 20 (5 years × 4 plots each). b) top 20 individual phylotypes, n = 20 (5 years × 4 plots each).

### Distribution of fungal lifestyles

Most phylotypes were either saprotrophs (soil: 119, litter: 71, wood: 62, unspecified: 29, other: 5), EMf (109, including 12 EM-borderline), or unassigned (343). Mycoparasites (11 phylotypes, 0.8% of all shares) comprised mainly *Trichoderma*. Further groups contained few taxa (e.g., sooty moulds: 1, root endophytes: 3) and/or were detected infrequently (e.g., plant pathogens: 17, 1 median read sample^-1^). Approximately half of the observations were of unknown lifestyles, with the rest distributed among EMf and saprotrophs (Fig. 6a).

**Figure 6:**
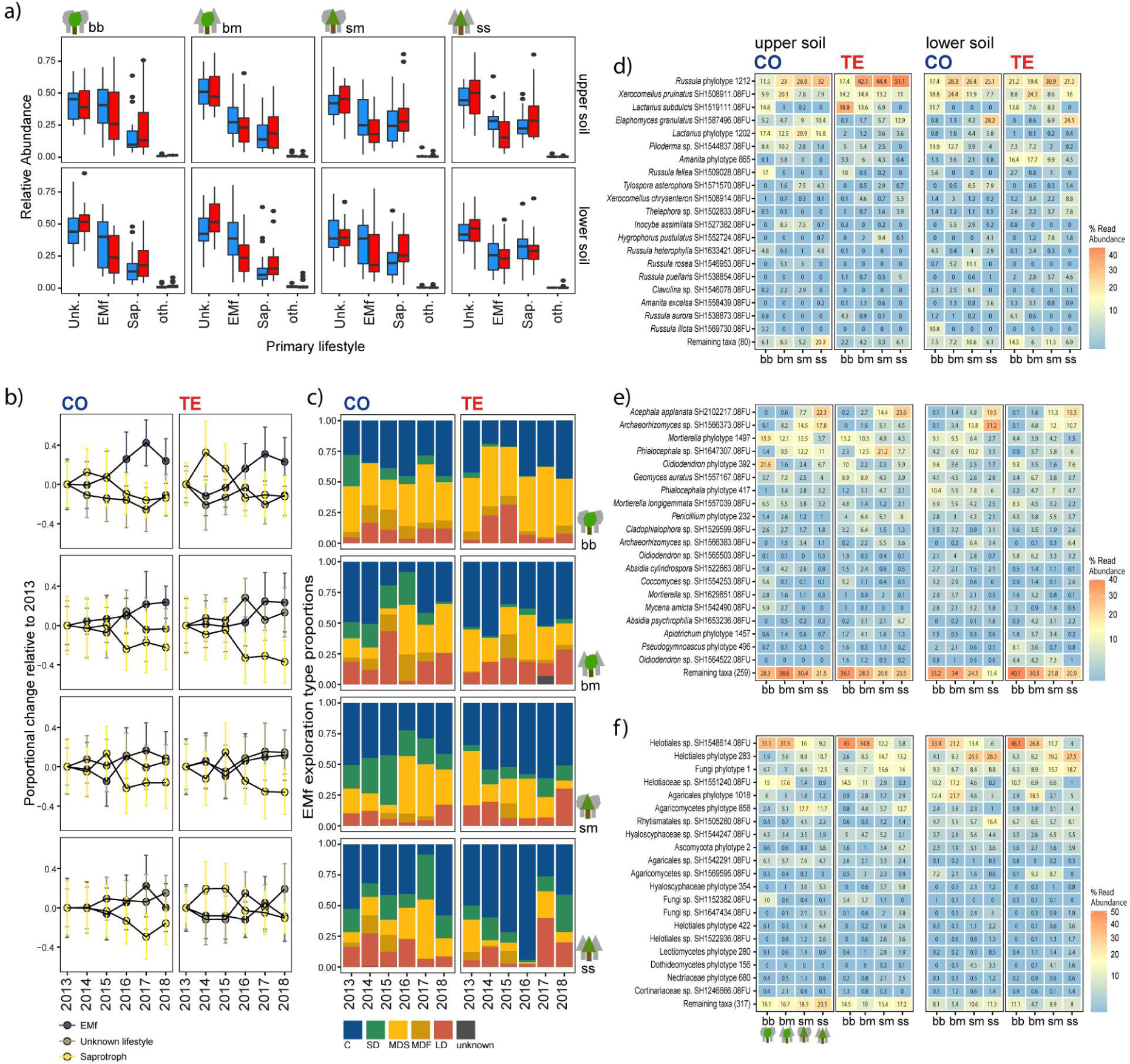
Distribution of fungal lifestyles. a) Overall assignments of primary fungal lifestyles during the experiment (2014-2018). Blue: CO, red: TE, Unk.: unknown lifestyle, EMf: ectomycorrhizal fungi, Sap.: saprotrophs, oth: ‘others’ including 15 different lifestyle groups (FunTraits primary lifestyle); n = 20 (5 years × 4 plots). b) Changes in relative proportions of major fungal lifestyles compared to their levels in 2013, i.e., before the start of the TE experiment (upper soil layer, lower soil in Fig. S3a); lilac: EMf, yellow: saprotrophs, grey: unknown lifestyles, n = 4, error-bars: 1 standard error of the difference in means respective to 2013. c) Ectomycorrhizal exploration types in the root-rhizosphere (upper soil layer, lower soil in Fig. S3b): Exploration types: C – contact, SD – short-distance, MDS – medium-distance-smooth, MDF – medium-distance-fringe, LD – long-distance, unknown – debated or unknown; d, e, f) TOP 20 most abundant phylotypes for EMf (d), saprotrophs (e), and fungi of unknown lifestyle (f); and their relative abundance within their group; n = 20 (5 years × 4 plots).

Different trends in shares of the major lifestyles were apparent during the earlier (2014-15) vs the later (2017-18) experimental years (Fig. 6b) with a juncture after the natural hot-drought summer of 2015: Before, saprotrophs tended to increase relative to EMf and unknowns. Afterwards, saprotrophs tended to remain stable or decline (depending on the root zone and soil layer) in TE and CO, while EMf tended to increase in all but spruce root zones in the upper (Fig. 6b) and lower soil (Fig. S3a).

### Individual distributions and temporal trends

The most abundant phylotypes for EMf, saprotrophs and unknown lifestyle are displayed in Fig. 6d-f. Yet, abundance patterns varied according to numerous factors (e.g., root zone, treatment, soil layer, legacy on the plots, sampling year), making an individual review of time courses (Figs. S4, S6, S7) necessary:

#### EMf

The most abundant among EMf (*Russula sp. 1212*) fluctuated. Nevertheless, the time series revealed a major peak in its relative share in spruce roots in 2016 TE and 2015 CO, especially when averaging soil layers (Fig. S4c). *Xerocomellus pruinatus* strongly varied without a consistent temporal pattern but had higher shares in lower soil. *Lactarius subdulcis,* absent in pure spruce, showed dissimilar trends in the other zones. In pure beech, its relative share increased from 2013 to 2017/2018, especially in TE, while it remained constant in TE beech mixture and strongly declined in TE spruce mixture from 2016-2018 (Fig. S4c). *Elaphomyces granulatus* fluctuated and showed no response towards TE, but strongly preferred lower soil of pure spruce where it dominated for three years. *Lactarius* sp. 1202 occurred in all root zones and preferred upper soil where it increased in CO in later years and strongly declined in TE after three years of treatment. Its decline was least in pure beech (6×), and biggest in pure spruce (43×) with mixture in-between. *Amanita sp. 865* increased with TE, especially in lower soil, where it became one of the most abundant EMf in later years, excepting pure spruce (Fig. S4b).

Many EMf were detected in amounts ≥ 0.1% infrequently or only under specific conditions, but patterns were often visible. For example, *Russula aurora* occurred regularly in lower soil and its share increased in TE pure beech over time compared to CO. Others were prominent in individual years, such as *Russula heterophylla,* which was highly abundant in CO 2018 but negligible earlier (Fig. S4c).

EMf exploration types mostly differed with root zones (Fig. 6c, Fig. S3b). They showed annual variation but remained similar between CO and TE with tendencies for more medium-distance types in TE beech and fewer short-distance types in TE spruce. An exception was upper soil TE pure spruce, where shares for contact types strongly increased until 2016 (Fig. 6c).

EMf functionality, displayed by extracellular enzyme activities (EA), was similar between treatments on the level of vital EM tips (EA_tip_) throughout the experiment for both tree species (Fig. S5a,b). Exceptions were significantly elevated EA_tip_ in TE pure spruce 2016 for carbohydrate-active enzymes (cellobiohydrolase, glucoronidase, β-glucosidase) and laccase that generally deviated from the other EA. EA extrapolation to soil volumes (EA_vol_) mirrored the numbers of vital tips (Fig. S5c,d), i.e., both showed similar patterns of reduced quantities.

#### saprotrophs

The most abundant, *Acephala applanata* and *Archaeorhizomyces* SH1566373.08FU, strongly preferred pure spruce (c. 11-31% of saprotrophs in pure spruce vs c. 0.1% in pure beech, Fig. 6e; Fig. S6c). SH1566373.08FU declined over time in TE spruce zones but remained stable in CO spruce (dominating lower soil). *A. applanata* became by far the most abundant saprotroph in upper soil TE pure spruce in later years (33-42% 2016-2018), where the other declined (Fig. S6a). *Mortierella sp*. was affine to upper soil beech, where it dominated in 2014 and 2015. *Phialocephala* SH1647307.08FU was common in pure spruce and mixtures where it rose from medium (2-8% upper soil 2013) to dominating saprotroph from 2016 onwards, especially in TE upper soil mixture (beech: 15-23% 2016-2018, spruce 28-43% 2016-2018). *Oidiodendron* sp. was especially common in monospecific zones of both treatments from 2016 onwards (Fig. S6).

#### ungi with unknown lifestyles

Half of the most common phylotypes with unknown lifestyles were assigned to species hypotheses, although taxonomical resolution varied similarly for all (Fig. 6f).

Helotiales were prominently represented, with SH1548614.08FU being the most abundant of all fungi (Fig. 5b). It dominated the unknown group in pure beech and to a lesser degree in beech mixture (on average >30% in upper soil) and occurred less towards pure spruce (averaging 9/4% upper/lower soil). The very abundant Helotiales phylotype 283 followed an opposing trend with higher abundances towards pure spruce. Another common Helotiaceae (SH1551240.08FU) was very affine to beech (upper soil), making up 11-18% in beech pure and mixed zones. Notably, spruce zones had more medium-abundant (1-10%) phylotypes of unknown lifestyle in upper soil compared to beech (Fig. S7a). Clear differences between CO and TE or decisive temporal trends were rare in the group (Fig. S7).

## Discussion

### Transient change in root associated fungi during five years TE

Overall, we did not find evidence for progressively changing α-diversity, community composition and functions (fungal lifestyles, EM extracellular EA) due to repeated drought. Instead, TE lead to a shift in root fungal communities that classify it as an environmental determinant similar to tree species’ root zone or soil layer. Such community shifts according to stress have also been observed for all soil fungal guilds at the treeline within a three years warming experiment (Solly et al., 2017).

Noteworthy, TE varied in importance according to fungal lifestyle. While TE was on par with the root zone for EMf composition depending on the year, it played a much smaller role for the unknown lifestyle group with saprotrophs in between. In soil fungal communities, EMf may respond less than saprotrophs (Castaño et al., 2018). However, stronger dependent free-living soil saprotrophs compared to those oriented to living at roots are to be expected in drying soils. Besides, this order of effect strength raises questions about the actual lifestyle of root fungi that are unknown to trait databases but also those defined as saprotrophs (cf. next chapters).

Additionally, TE effect sizes varied during the time course, strongest for EMf and least for unknowns. Hereby, the biggest community shifts relative to CO were seen after three years TE for all major functional groups, despite continued low soil water contents during the later years. This aligns with pre-dawn leaf water potentials (Grams et al., 2021), indicating that the trees were most stressed up to year three of the experiment. It is possible that tree acclimation to long-term low soil moisture (growth reductions above and belowground (Grams et al., 2021)) relaxed the drought stress by reducing leaf area and thus water demand by transpiration (Gebhardt et al., 2023). As very subtle improvements in the soil water status were reported to sustain fine root growth under otherwise dry conditions (Joseph et al. 2020), we may also see acclimation of the tree-soil system to long-term drought.

Taken together, this suggests that TE over multiple growth periods in beech and spruce forests can deteriorate root fungal communities, but stress on the host trees needs to be almost fatal (above the 69 % precipitation reduction in our experiment (Grams et al., 2021)) otherwise fungal communities may just shift to an altered stable state recognisably similar to CO. Supporting this, a lighter precipitation reduction (25%) in a *Quercus ilex* forest did not even change fungal communities after five years (Bastida et al., 2019).

We therefore reject H1 that extended drought would progressively alter root associated fungal communities but confirm that it leads to a shifted composition that may still deteriorate depending on the drought stress of their host trees.

### Importance of vital fine roots for the biotrophic subset

For EMf, the number of vital root tips mattered when they became extremely low. Earlier results during the first three years TE showed a progressively decreasing diversity of spruce EMf and increasingly diverging EMf compositions, while EA remained comparable between CO and TE (Nickel et al. 2018). Here, we found a contact exploration type (*R. ochroleuca* according to personal observations on fruiting bodies and morphotyping (SI Nickel et al., 2018)) increasingly and extraordinarily dominating in TE spruce up to the third year, indicating that the distinct loss of vital tips for spruce resulted in a highly disturbed system. *R. ochroleuca* may have been most efficient in colonising reactivated dormant root tips, possibly favoured by rhizomorphs that grow along roots (Agerer, 2006). Speculatively, we observed a rapid form of drought-related community disassembly towards generalists, which Gehring et al. (2014) found in a long-term study on *Pinus edulis*.

Adverse to spruce, strongly diminished fine-root numbers did not lead to signs of deterioration in beech fungal communities, likely due to maintained fine-root production (Zwetsloot & Bauerle, 2021), which prevented a local loss of habitats.

While mirrored in results on root production (Zwetsloot and Bauerle, 2022), we did not expect the partial recovery of fine roots in 2017-2018, which was paralleled by recovered spruce EMf diversity, less divergent fungal communities, and less stressed trees (see above). Substantial contributions by the EMf community to relaxing tree stress seem unlikely, especially as long-distance exploration types did not profit from TE in later years, questioning their advantages under long-term drought for mature trees (Letho and Zwiaczek, 2011; Nickel et al., 2018; Castaño et al., 2023). Adversely, the extremely dry TE soil may have led to hydraulic redistribution of water from trees to EMf, which could be vital for the nutrient support of the host trees but remains rarely studied (cf. Querejeta et al., 2003; Letho and Zwiaczek, 2011). Castaño et al. (2018) observed shifts from long- to short-distance EMf coupled with decreasing free living saprotroph during natural drought, which can be interpreted as an effect of a soil moisture gradient towards roots supported by hydraulic redistribution.

Although the EMf diversity overall did not deteriorate, clear tree species and root-zone-specific reactions of major EMf were seen. Two highly abundant *Lactarius* phylotypes exemplify that drought reactions can be strongly modulated by the tree host. *L. subdulcis* profited from TE in pure beech and declined in pure spruce compared to CO. In the TE plots, beech litter accumulated (Brunn et al., 2023), which could have benefitted this fungus with a preference for root colonisation in slightly decomposed beech litter (personal observation). The other *Lactarius* phylotype was immediately reduced to minor amounts by TE and could not recover later.

Though database-designated endophytes and pathogens were negligible, many endophytes likely remained unrecognised within the unknown-lifestyle group. Clear indications are a remarkably strong and stable dependence of the unknowns on their respective root zone and a weak response to TE that only became apparent when the trees were most stressed. Nonetheless, high numbers of unresolved taxa were still expected (Tanunchai et al., 2023).

In summary, evidence was limited for progressively changing biotrophic fungi with TE (H2). Instead, TE-EMf remained significantly shifted but remarkably related to CO, while a group of putative endophytes was barely influenced. Biotrophic root fungal communities at the root tips of TE presented themselves as surviving islands potentially dependent on tree-supplied water through hydraulic redistribution. However, different fine root growth strategies between beech and spruce likely make the latter more prone to EMf community deterioration during extreme drought (H2a).

### Root saprotrophs responded differentially to reduced moisture

The response of saprotrophs to TE was between those of EMf and unknowns, and likely based on abundant subgroups reacting individually.

Our findings *on Archaeorhizomyces* agree with reports designating them as a substantial component of the rhizosphere (Menkis et al., 2014) with tree species specificity and preference to lower soil (Pinto-Figueroa et al., 2019). Losses through TE align with results from a spatiotemporal gradient in Switzerland, where *Archaeorhizomyces* preferred colder, more humid conditions (Pinto-Figueroa et al., 2019). Although their ecological role remains unclear, they are potential keystone taxa (Choma et al., 2016) whose drought sensitivity may leave a gap in the spruce root-fungal system.

Dark septate endophytes (DSE), here mainly *Acephala* and *Phialocephala* complexes, increased during TE. Consistently, *A. applanata* preferred drier conditions in mature oak (Landolt et al., 2020) and spruce seedlings (Stroheker et al., 2018) but also increased with dead root material in opposition to other DSE (Stroheker et al., 2018). Their role in our experiment remains unsolved as their lifestyle can range from mutualism to parasitism (cf. Landolt et al., 2019; Terhonen, 2021). Yet, DSE can improve plant function under harsh conditions and increase in water-limited environments (cf. Terhonen, 2021). Thus, the increase of DSE in CO after 2015 may indicate that the extreme natural drought of that year gave them an advantage, for example by protecting their host roots from drying out (Landolt et al., 2020), and they were able to maintain stable abundance thereafter without exceptional stress (Terhonen, 2021).

*Mortierella* comprising ubiquitous and versatile saprotrophs (Nagy et al., 2011; Uehling et al., 2017) likely functioned as early-stage decomposers in the upper soil of CO and TE until 2015, and got partly replaced by *Oidiodendron* later, a taxon with broad mutualistic capabilities (Martino et al., 2018), indicating a shift to root interaction rather than decay.

Interestingly, mineral nitrogen in upper soil shifted from lower levels until 2015 to higher levels afterwards (in both, CO and TE), suggesting that more nitrogen had been released than could be used later. While we did not assess decay rates, soil organic matter had accumulated on TE compared to CO at the end (Brunn et al., 2023), which shows that drought strongly affected decay rates in our system. The relative increase of saprotrophs during the early phase of TE pure beech and spruce could have been due to accumulating dead roots, which resulted in primed fungal decomposition (Kuzyakov, 2010; Zhou et al., 2021). Although we expected an increasing effect of TE over time, our results rather suggest a prolonged impact of the 2015 natural drought and/or an acclimation effect of the belowground system to the long-term drought.

Time-courses for saprotrophs were much less concentrated on few highly abundant phylotypes than for EMf but showed a wide range of high to medium-abundant saprotrophs and many rare ones, especially for beech, suggesting that the continuously renewing fine roots of beech are colonised rather according to stochastic selection (cf. Zhou & Ning, 2017) of root-associated saprotrophs from the soil. This is also supported by Bahram et al. (2016), who found saprotrophic fungi among the most stochastically selected Eukarya in a boreal forest.

Overall, effects of low soil moisture on saprotrophs were found with TE (H3), though not coupled with reduced diversity but community shifts towards fungi with stronger biotrophic capabilities.

### Tree mixture effects

Tree mixture had no discernible effect on the fluctuating numbers of vital root tips in our experiment (a positive impact on spruce roots remains possible, according to Zweetslot & Bauerle, 2022). Furthermore, the fungal α-diversity of two tree species in pure root zones was not added to the roots of a single species in the mixed zone. Similarily, EMf richness did not increase with more tree species in a boreal system (Otsing et al., 2021). Likewise, Tedersoo et al. (2016) found that soil microbial diversity may depend rather on the general soil context than on tree species numbers. Supporting this, soil microbial communities in mixtures of beech and spruce were found to be hybrids of the respective pure stands (Wilhelm et al., 2023).

However, tree mixture improved fungal communities under TE by attenuating most high abundance taxa compared to pure root zones, thus carrying the potential to add functional redundancy that could be rather limited in forest soil fungi (Mori et al., 2016).

While a tree mixture effect on community composition would be expected in soil due to the changed litter quality (e.g., Aponte et al., 2010; Yang et al., 2022), it is remarkable that root fungal sub-communities of all major lifestyles from mixture lay in between their pure zone counterparts. This highlights that tree root community compositions depend on neighbouring tree species, as seen for spruce EMf by Otsing et al. (2021).

In summary, H4 (niche diversification through tree mixing creates an advantage under TE) can be cautiously accepted under the precondition that the stress needs to be severe. However, fungal diversity is not higher in TE mixture compared to CO (rejecting H4a). Instead, influences of single taxa are attenuated in CO and TE, while a more efficient soil exploration in TE mixture is not deducible from our exploration type results (rejecting H4b). Actual benefits of attenuated dominating taxa need to be carved out in future studies.

### Conclusion

Despite significant differences, the overall similarity between TE and CO fungal communities and their functions after five drought years highlights the qualitative resilience of the fine root-fungal system. On the one hand, this fits very well with results that quantitative effects (e.g., fewer leaves – fewer fine roots) may have governed tree acclimation. On the other hand, it raises the question if the surviving vital tips, altogether a root-fungal system, may function as moist islands within dried-out soil, kept alive by an interplay between tree-redistributed water and fungal symbionts (e.g., prevention of water loss through DSE, nutrient acquisition by EMf). Finding out about these is a challenging but important goal for future studies on forests increasingly threatened by weather extremes in a changing climate.

## Supporting information

Collection of all supplementary data V1.0

## Acknowledgements

This work was funded by the German Research Foundation (DFG) through grant MU 831/23-1 and DFG PR555/2-1, and by the Bavarian State Ministries of the Environment and Consumer Protection as well as Food, Agriculture and Forestry (BayKROOF, W047/Kroof II).

We are grateful to Bärbel Groß, Elke Gerstner, and Franz Buegger for their invaluable help in sampling and sample processing over many years, and to our volunteers in ecology Laura Pohlenz and Valentin Kugler for their assistance.

## Author contributions

FW and KP designed the experiment. FW processed the samples and analysed the data. FW, TG, and KP interpreted the data. FW and KP wrote the manuscript with contributions from TG.

## Data availability

All raw sequencing data connected to this study are openly available in in the NCBI Sequence Read Archive under the BioProject accession number PRJNA985992. All other data presented in this study are available in this article and the respective Supplementary Materials.

The following Supporting Information is available for this article:

**Fig. S1** Soluble inorganic nitrogen contents

**Fig. S2** Diversity of all fungi and major lifestyle groups

**Fig. S3** Changes in relative proportion of fungal lifestyles (lower soil)

**Fig. S4** Time-courses of EMf

**Fig. S5** EM extracellular enzyme activities

**Fig. S6** Time-courses of saprotrophs

**Fig. S7** Time-courses of fungi with unknown lifestyle

**Table S1** Primer sequences

**Table S2** Annual PERMANOVA results with reduced experimental units

**Table S3** PERMANOVA results for Fig. 4 including interactions

**Methods S1** Ectomycorrhizal enzyme activity tests

