## Supplementary material for "Responses of root-associated fungal communities of mature beech and spruce during five years of experimental drought": Collection of all supplementary data V1.0

The following Supporting Information is available for this article:

**Fig. S1** Soluble inorganic nitrogen contents

**Fig. S2** Diversity of all fungi and major lifestyle groups

**Fig. S3** Changes in relative proportion of fungal lifestyles and EMf exploration types (lower soil)

**Fig. S4** Time-courses of EMf

**Fig. S5** EM extracellular enzyme activities

**Fig. S6** Time-courses of saprotrophs

**Fig. S7** Time-courses of fungi with unknown lifestyle

**Table S1** Primer sequences

**Table S2** Annual PERMANOVA results with reduced experimental units

**Table S3** PERMANOVA results for Fig. 4 including interactions

**Methods S1** Ectomycorrhizal enzyme activity tests

**Fig. S1** Soluble inorganic nitrogen compounds after $CaCl_2$-extraction from control (CO, blue) and drought plots (TE, red) in the upper (dashed lines) and lower (dotted lines) soil layer during the period of throughfall exclusion. Error bars: ±1 standard error. * and °: Wilcoxon $p < 0.05$ (CO vs. TE), * for upper soil, ° for lower soil; $n_{units}$ = 4-6 (two experimental units removed after 2015 due to bark beetle attack).

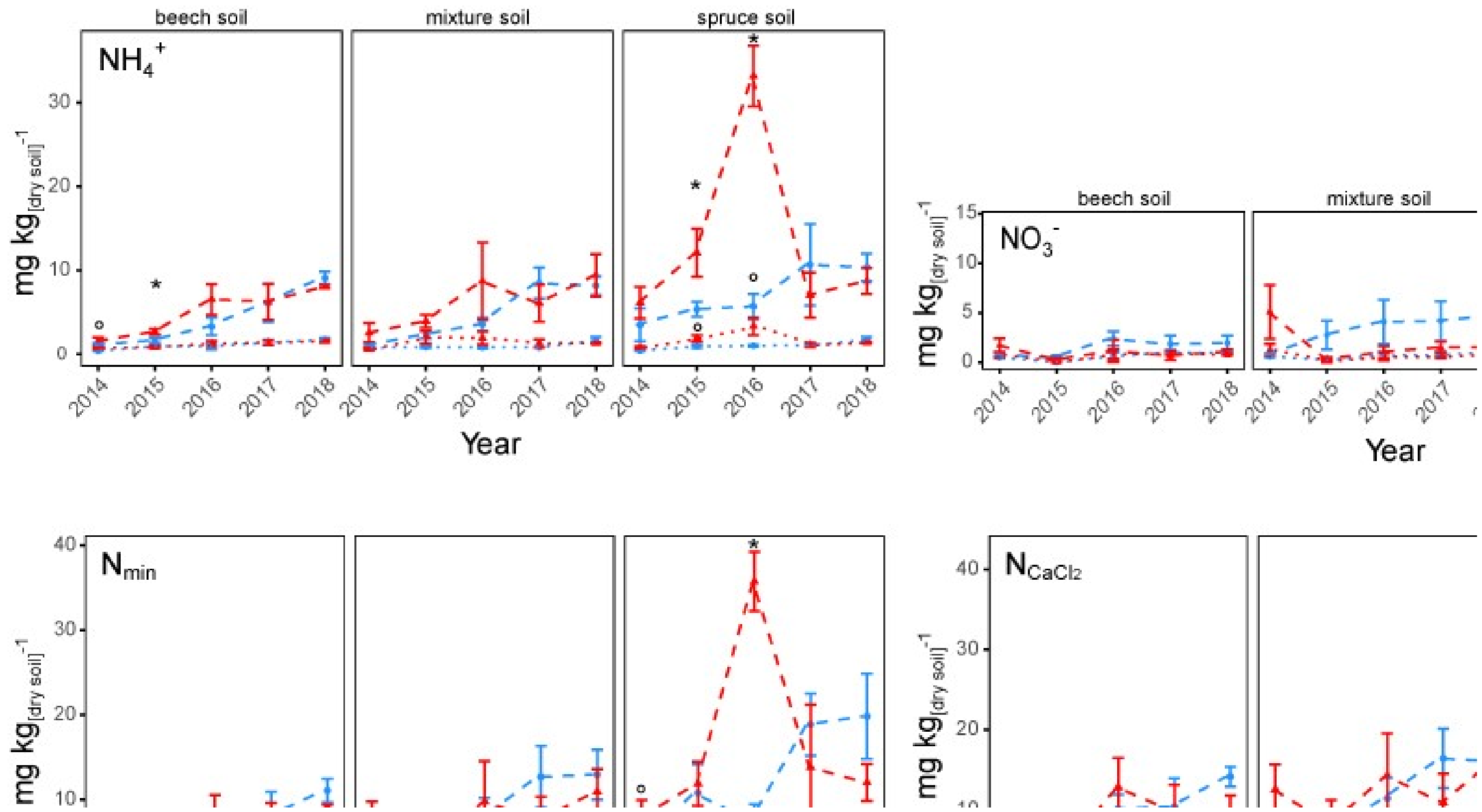

**Fig. S2** Diversity indices for the total fungal community (a), and Simspon Index (b) as well as Evenness (c) for major lifestyle groups in controls (CO, blue) and drought treatment (TE, red) plots. Lines in (b,c): solid = ectomycorrhizal fungi, dashed = fungi of unknown lifestyle, dotted = saprotrophs; bb, bm, sm, ss: different root zones (bb – pure beech, bm – beech mixture, sm – spruce mixture, ss – pure spruce); error bars: ±1 standard error.

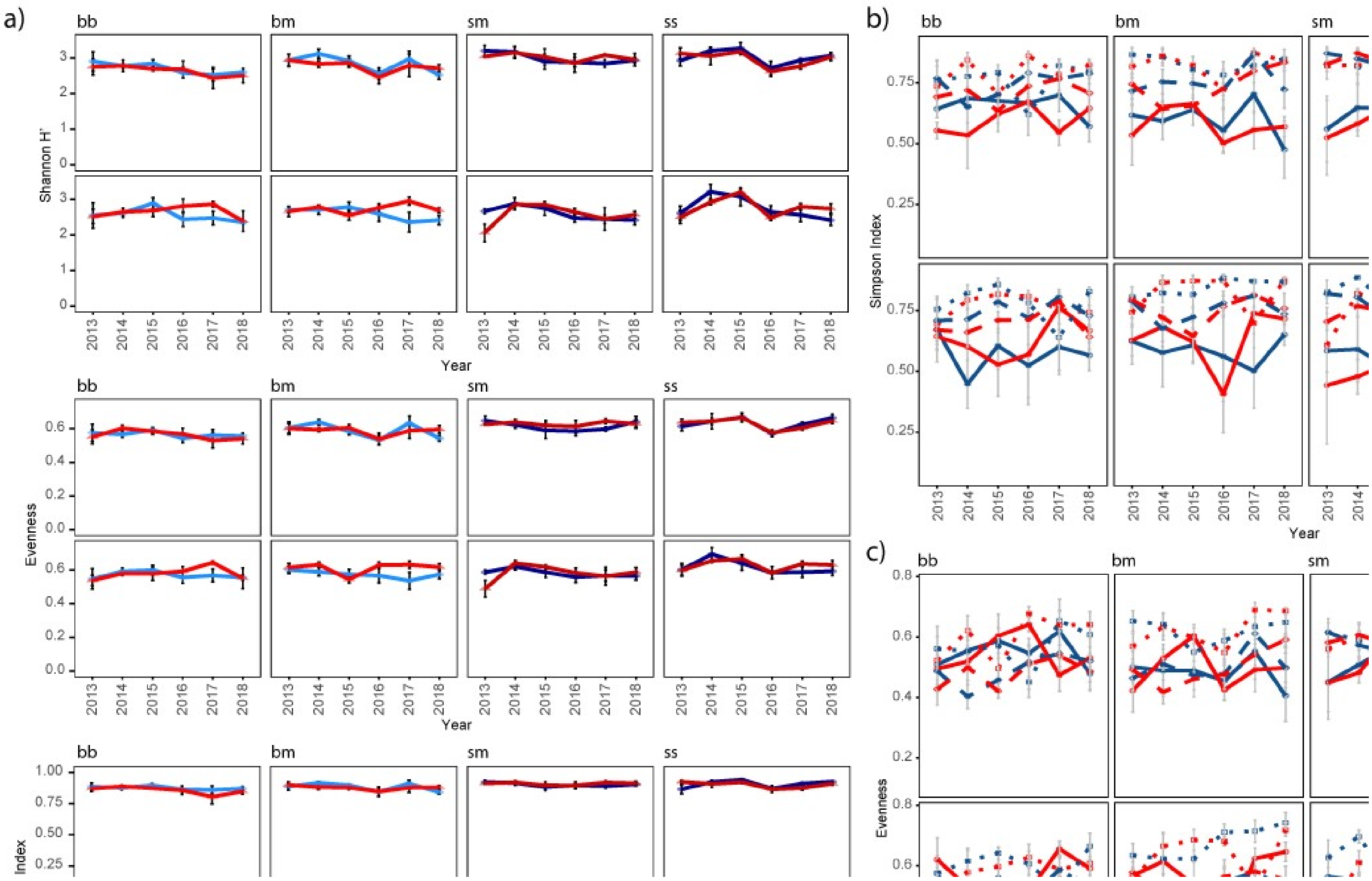

**Fig. S3** Lower soil layer: changes in relative proportion of major fungal lifestyles compared to their levels in 2013 (before the start of the TE experiment) (a) and ectomycorrhizal fungal (EMf) exploration types in the root-rhizosphere (b); $n_{units}$ = 4, error-bars in (a): 1 standard error of the difference in means respective to 2013. CO: control plots, TE throughfall exclusion plots; bb, bm, sm, ss: different root zones (bb – pure beech, bm – beech mixture, sm – spruce mixture, ss – pure spruce); exploration types: C – contact, SD – short-distance, MDS – medium-distance-smooth, MDF – medium-distance-fringe, LD – long-distance, unknown – debated or unknown

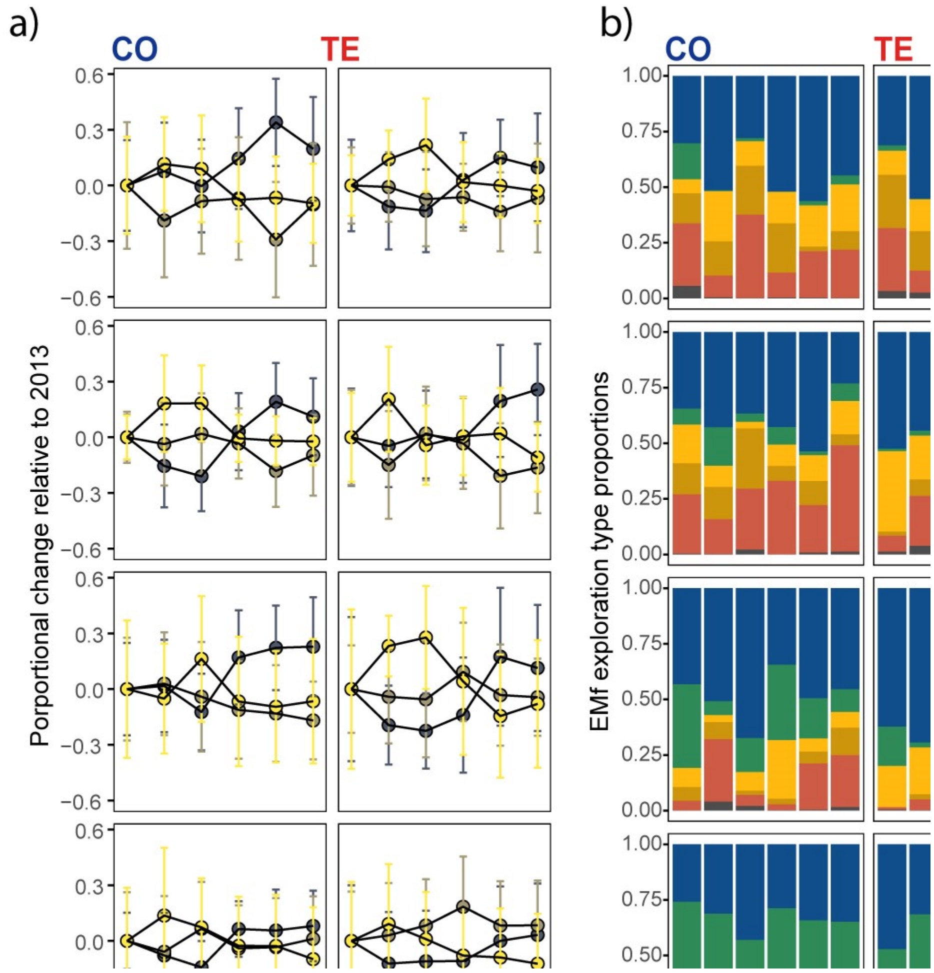

**Fig. S4** Heatmap time-courses for EMf fungi in upper (S4a) and lower soil layers (S4b) and averaged for both layers (S4c); bb, bm, sm, ss: root zones (bb – pure beech, bm – beech mixture, sm – spruce mixture, ss – pure spruce); CO: control plots, TE: throughfall exclusion plots

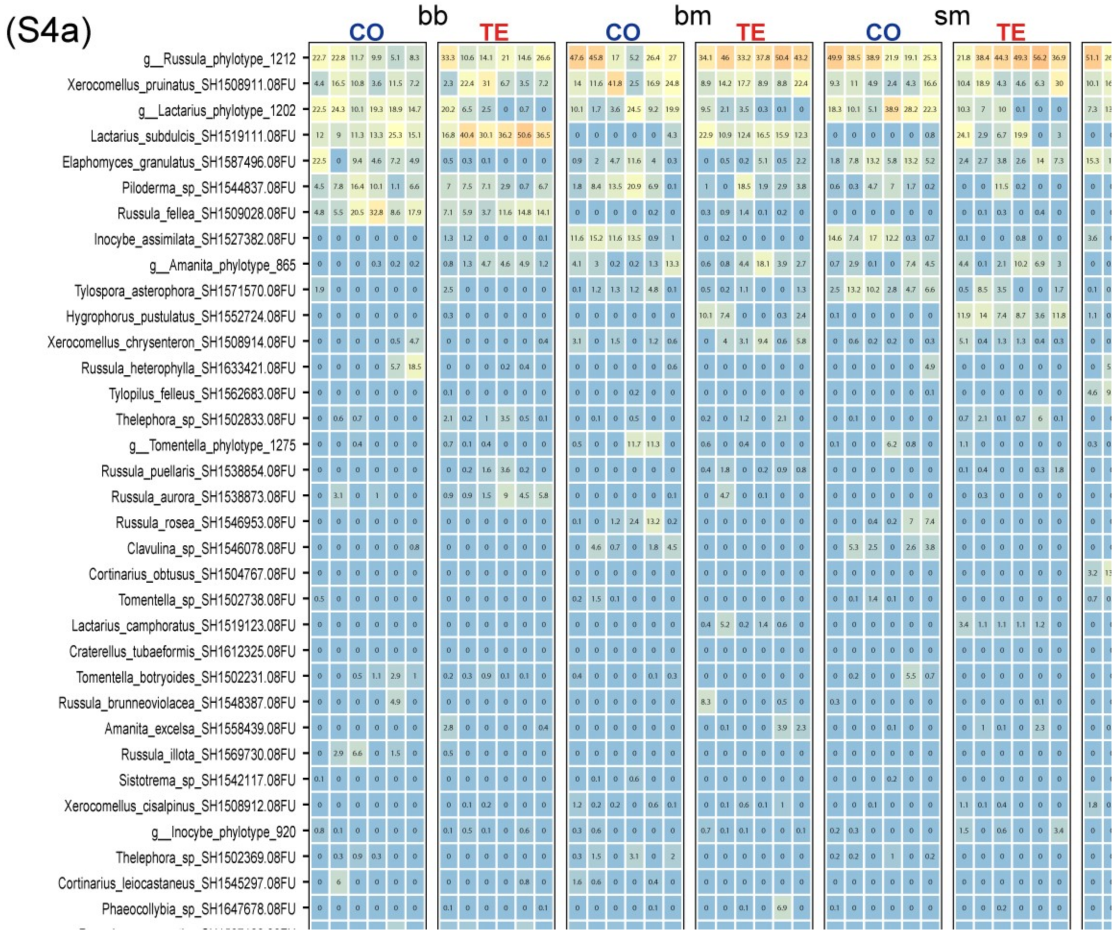

(S4b)

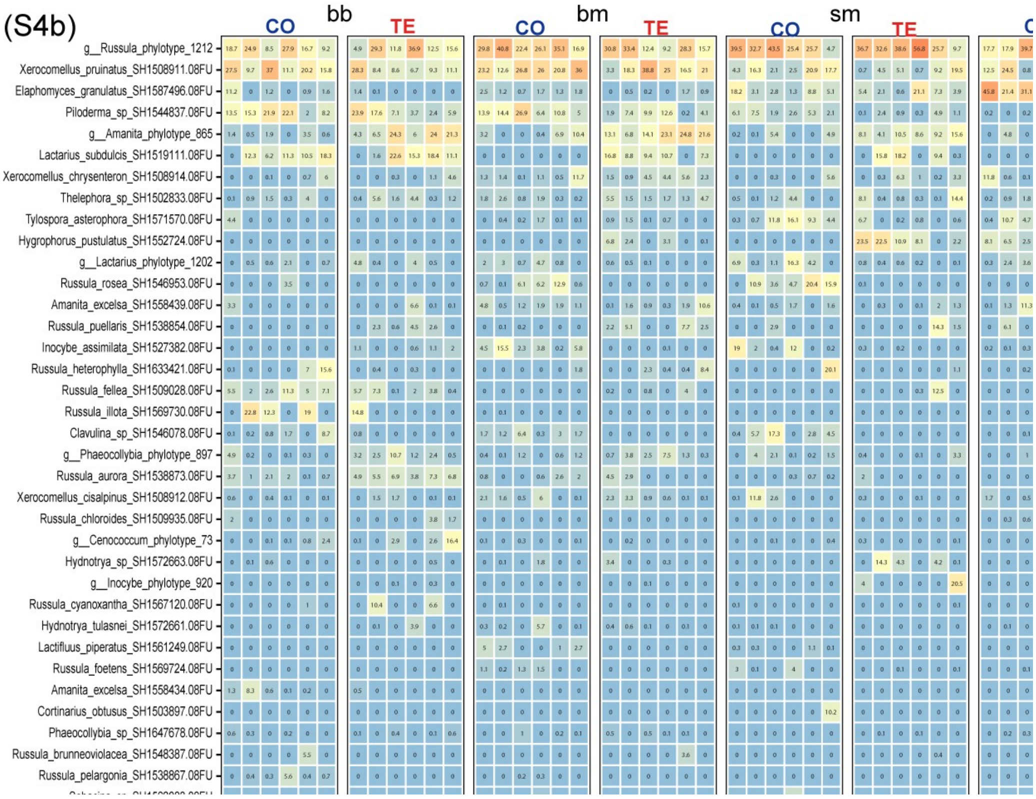
bb
bm
sm
CO
TE
CO
TE
CO
TE
g__Russula_phylotype_1212
Xerocomellus_pruinatus_SH1508911.08FU
Elaphomyces_granulatus_SH1587496.08FU
Piloderma_sp_SH1544837.08FU
g__Amanita_phylotype_865
Lactarius_subdulcis_SH1519111.08FU
Xerocomellus_chrysenteron_SH1508914.08FU
Thelephora_sp_SH1502833.08FU
Tylospora_asterophora_SH1571570.08FU
Hygrophorus_pustulatus_SH1552724.08FU
g__Lactarius_phylotype_1202
Russula_rosea_SH1546953.08FU
Amanita_excelsa_SH1558439.08FU
Russula_puellaris_SH1538854.08FU
Inocybe_assimilata_SH1527382.08FU
Russula_heterophylla_SH1633421.08FU
Russula_fellea_SH1509028.08FU
Russula_illota_SH1569730.08FU
Clavulina_sp_SH1546078.08FU
g__Phaeocollybia_phylotype_897
Russula_aurora_SH1538873.08FU
Xerocomellus_cisalpinus_SH1508912.08FU
Russula_chloroides_SH1509935.08FU
g__Cenococcum_phylotype_73
Hydnotrya_sp_SH1572663.08FU
g__Inocybe_phylotype_920
Russula_cyanoxantha_SH1567120.08FU
Hydnotrya_tulasnei_SH1572661.08FU
Lactifluus_piperatus_SH1561249.08FU
Russula_foetens_SH1569724.08FU
Amanita_excelsa_SH1558434.08FU
Cortinarius_obtusus_SH1503897.08FU
Phaeocollybia_sp_SH1647678.08FU
Russula_brunneoviolacea_SH1548387.08FU
Russula_pelargonia_SH1538867.08FU

(S4c)

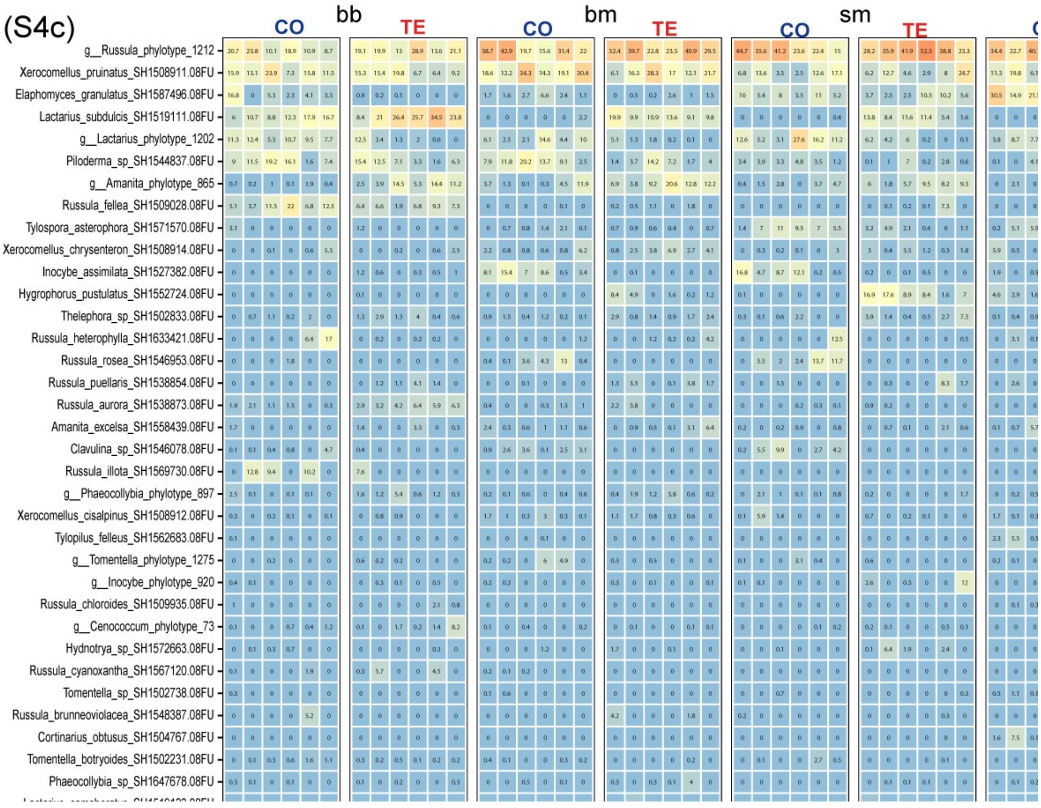
bb
bm
sm
CO
TE
CO
TE
CO
TE
g__Russula_phylotype_1212
Xerocomellus_pruinatus_SH1508911.08FU
Elaphomyces_granulatus_SH1587496.08FU
Lactarius_subdulcis_SH1519111.08FU
g__Lactarius_phylotype_1202
Piloderma_sp_SH1544837.08FU
g__Amanita_phylotype_865
Russula_fellea_SH1509028.08FU
Tylospora_asterophora_SH1571570.08FU
Xerocomellus_chrysenteron_SH1508914.08FU
Inocybe_assimilata_SH1527382.08FU
Hygrophorus_pustulatus_SH1552724.08FU
Thelephora_sp_SH1502833.08FU
Russula_heterophylla_SH1633421.08FU
Russula_rosea_SH1546953.08FU
Russula_puellaris_SH1538854.08FU
Russula_aurora_SH1538873.08FU
Amanita_excelsa_SH1558439.08FU
Clavulina_sp_SH1546078.08FU
Russula_illota_SH1569730.08FU
g__Phaeocollybia_phylotype_897
Xerocomellus_cisalpinus_SH1508912.08FU
Tylopilus_felleus_SH1562683.08FU
g__Tomentella_phylotype_1275
g__Inocybe_phylotype_920
Russula_chloroides_SH1509935.08FU
g__Cenococcum_phylotype_73
Hydnotrya_sp_SH1572663.08FU
Russula_cyanoxantha_SH1567120.08FU
Tomentella_sp_SH1502738.08FU
Russula_brunneoviolacea_SH1548387.08FU
Cortinarius_obtusus_SH1504767.08FU
Tomentella_botryoides_SH1502231.08FU
Phaeocollybia_sp_SH1647678.08FU

**Fig. S5** Potential enzyme activities (EAs) on the level of vital root tips ($EA_{tip}$) (a,b) and extrapolated to soil volumes ($EA_{vol}$) (c,d), each displayed as deviation from control (a,c) and alternatively with absolute values (upper soil only) (b,d). $EA_{tip}$ (pmol $cm^{-2}$ $min^{-1}$) is the weighted mean of EA per ectomycorrhizal (EM) tip in an EM fungal community (see method S1) and $EA_{vol}$ (pmol $cm^{-2}$ $min^{-1}$ $cm^{-3}$) is additionally taking into account the number of vital EM tips per soil volume. The layout for the graphs in (a) and (c) follows Nickel et. al. (2018): the grey dashed line with a slope of 1 and an intercept of 0 was drawn to indicate when EAs under control is equal to EAs under throughfall exclusion; deviation of the slope of regression lines from 1 with an intercept remaining close to 0 indicates similar relative degrees and directions of change in all EAs, whereas shift in the intercept indicates that EA values changed to different degrees and/or directions; values of intercept and slope are given in the top left corner of each panel with asterisks indicating significant differences from a slope of 1 and an intercept of 0 (*p < .05, **p < .01, ***p < .001); symbols represent sample types (circles pure beech root zone, squares mixed root zone of beech or spruce, triangles pure spruce) resulting in four values per enzyme and a total of 28 values per year of seven hydrolytic enzymes (Xyl - xylosidase, Cel-cellobiohydrolase, Gls - β-glucosidase, Nag - N-acetyl-glucosaminidase, Leu - leucine aminopeptidase, Pho - phosphatase and Glr - glucuronidase (laccase (Lac) is only shown in (b,d)). Error bars in (b,d): ±1 standard error. *: $p < 0.05$ (for CO vs. TE in a pairwise comparison tested using the function emmeans_test() (https://github.com/kassambara/rstatix) with bonferroni correction for multiple testing following a three-way ANOVA with the factors treatment, root zone, sampling year and random effect experimental unit); bb, bm, sm, ss: different root zones (bb – pure beech, bm – beech mixture, sm – spruce mixture, ss – pure spruce).

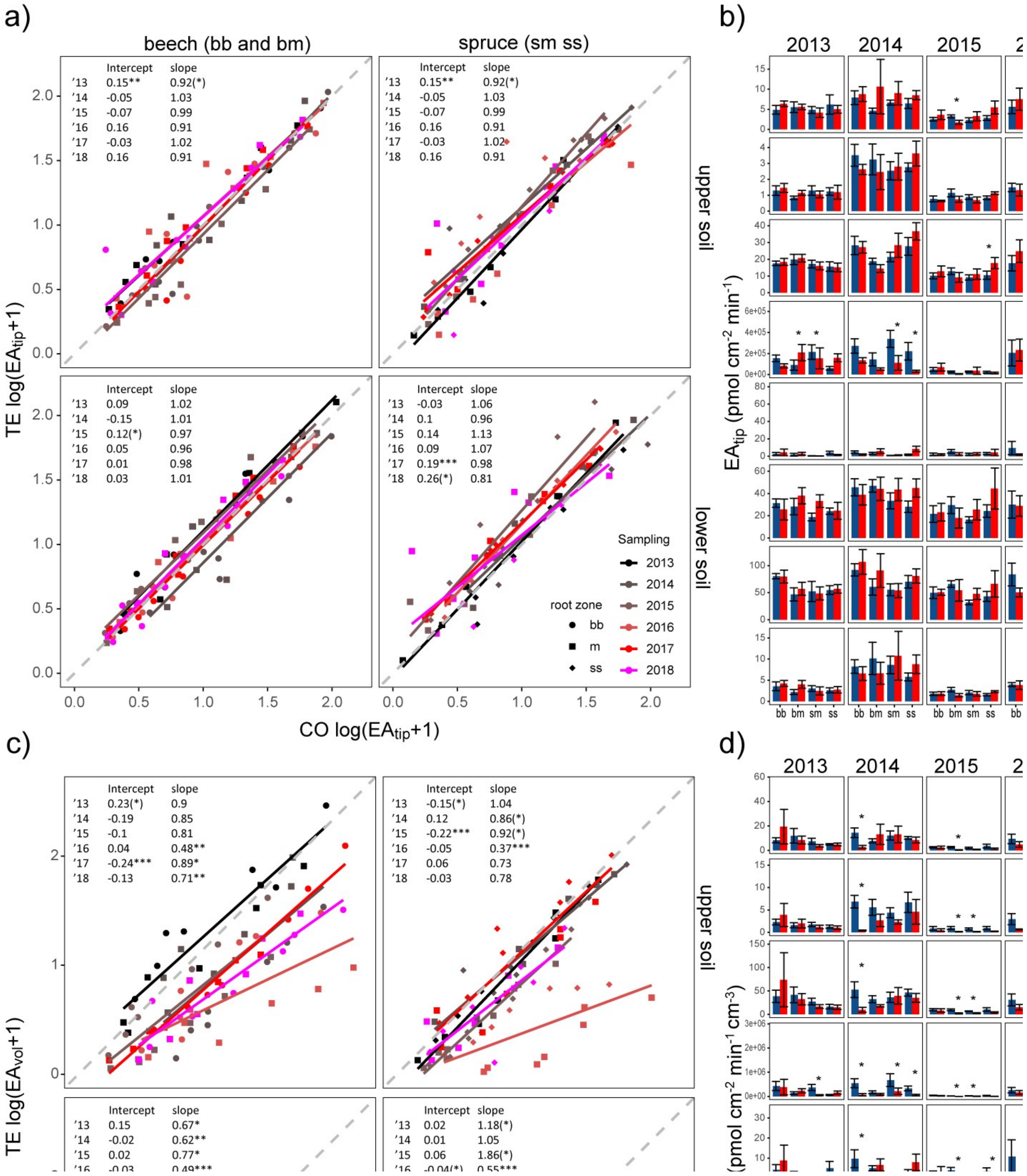
a)
beech (bb and bm)
spruce (sm ss)
Intercept slope
'13 0.15** 0.92(*)
'14 -0.05 1.03
'15 -0.07 0.99
'16 0.16 0.91
'17 -0.03 1.02
'18 0.16 0.91
Intercept slope
'13 0.15** 0.92(*)
'14 -0.05 1.03
'15 -0.07 0.99
'16 0.16 0.91
'17 -0.03 1.02
'18 0.16 0.91
upper soil
Intercept slope
'13 0.09 1.02
'14 -0.15 1.01
'15 0.12(*) 0.97
'16 0.05 0.96
'17 0.01 0.98
'18 0.03 1.01
Intercept slope
'13 -0.03 1.06
'14 0.1 0.96
'15 0.14 1.13
'16 0.09 1.07
'17 0.19*** 0.98
'18 0.26(*) 0.81
lower soil
Sampling
2013
2014
2015
2016
2017
2018
root zone
bb
m
ss
TE log(EA_tip+1)
CO log(EA_tip+1)
b)
2013
2014
2015
EA_tip (pmol cm^-2 min^-1)
bb bm sm ss
c)
Intercept slope
'13 0.23(*) 0.9
'14 -0.19 0.85
'15 -0.1 0.81
'16 0.04 0.48**
'17 -0.24*** 0.89*
'18 -0.13 0.71**
Intercept slope
'13 -0.15(*) 1.04
'14 0.12 0.86(*)
'15 -0.22*** 0.92(*)
'16 -0.05 0.37***
'17 0.06 0.73
'18 -0.03 0.78
upper soil
Intercept slope
'13 0.15 0.67*
'14 -0.02 0.62**
'15 0.02 0.77*
Intercept slope
'13 0.02 1.18(*)
'14 0.01 1.05
'15 0.06 1.86(*)
TE log(EA_vol+1)
d)
2013
2014
2015
(pmol cm^-2 min^-1 cm^-3)

**Fig. S6** heatmap time-courses for saprotrophic fungi in upper (S6a) and lower soil layers (S6b) and averaged for both layers (S6c); bb, bm, sm, ss: root zones (bb – pure beech, bm – beech mixture, sm – spruce mixture, ss – pure spruce); CO: control plots, TE: throughfall exclusion plots

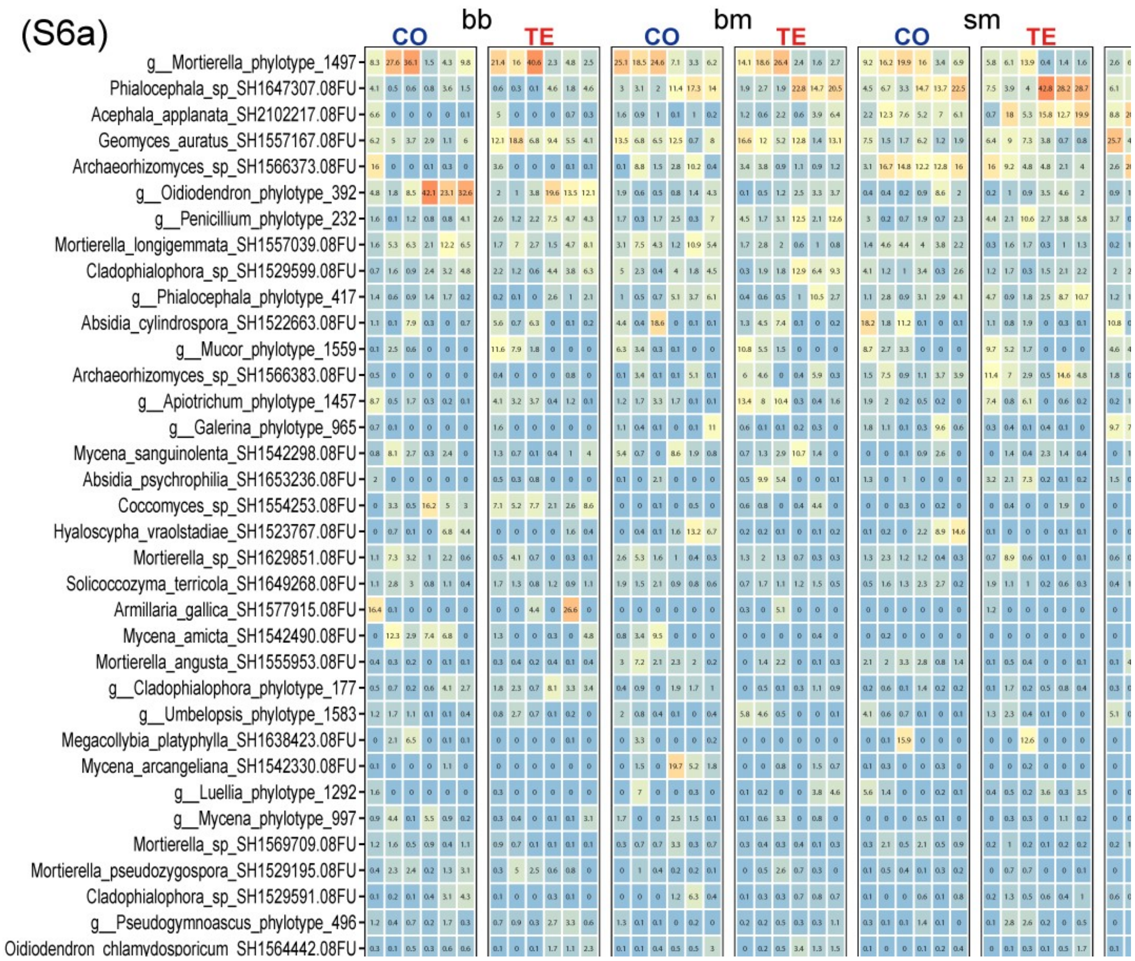

(S6b)

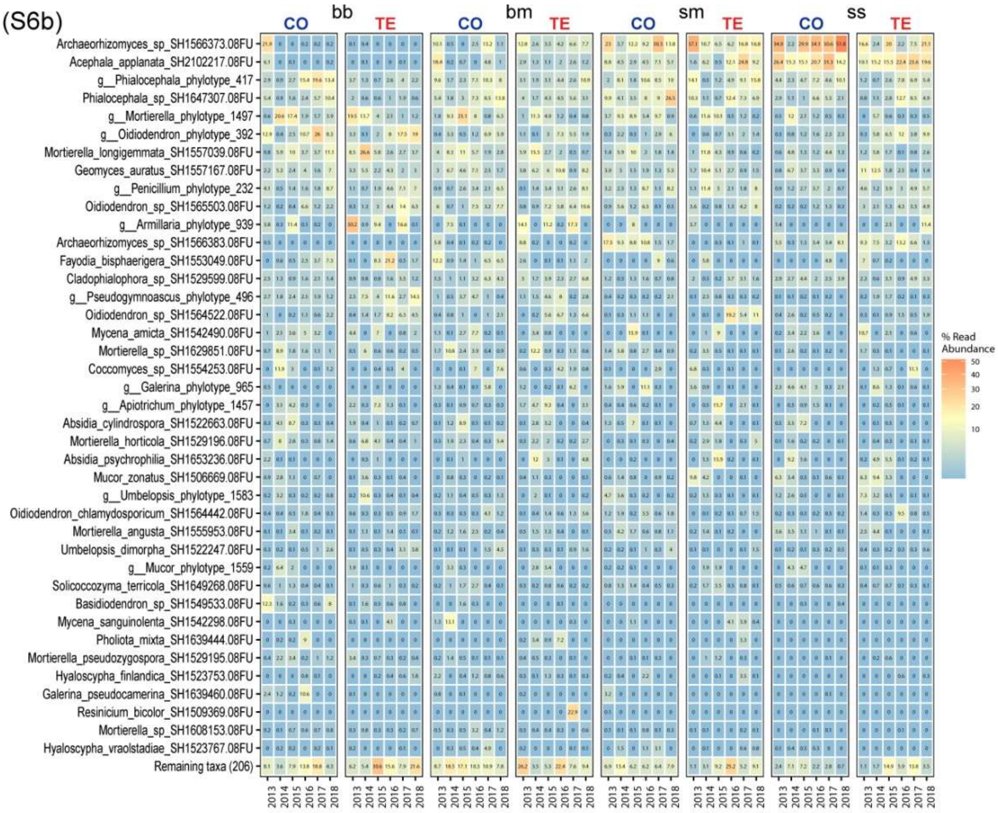

bb
bm
sm
ss
CO
TE
Archaeorhizomyces_sp_SH1566373.08FU
Acephala_applanata_SH2102217.08FU
g__Phialocephala_phylotype_417
Phialocephala_sp_SH1647307.08FU
g__Mortierella_phylotype_1497
g__Oidiodendron_phylotype_392
Mortierella_longigemmata_SH1557039.08FU
Geomyces_auratus_SH1557167.08FU
g__Penicillium_phylotype_232
Oidiodendron_sp_SH1565503.08FU
g__Armillaria_phylotype_939
Archaeorhizomyces_sp_SH1566383.08FU
Fayodia_bisphaerigera_SH1553049.08FU
Cladophialophora_sp_SH1529599.08FU
g__Pseudogymnoascus_phylotype_496
Oidiodendron_sp_SH1564522.08FU
Mycena_amicta_SH1542490.08FU
Mortierella_sp_SH1629851.08FU
Coccomyces_sp_SH1554253.08FU
g__Galerina_phylotype_965
g__Apiotrichum_phylotype_1457
Absidia_cylindrospora_SH1522663.08FU
Mortierella_horticola_SH1529196.08FU
Absidia_psychrophilia_SH1653236.08FU
Mucor_zonatus_SH1506669.08FU
g__Umbelopsis_phylotype_1583
Oidiodendron_chlamydosporicum_SH1564442.08FU
Mortierella_angusta_SH1555953.08FU
Umbelopsis_dimorpha_SH1522247.08FU
g__Mucor_phylotype_1559
Solicoccozyma_terricola_SH1649268.08FU
Basidiodendron_sp_SH1549533.08FU
Mycena_sanguinolenta_SH1542298.08FU
Pholiota_mixta_SH1639444.08FU
Mortierella_pseudozygospora_SH1529195.08FU
Hyaloscypha_finlandica_SH1523753.08FU
Galerina_pseudocamerina_SH1639460.08FU
Resinicium_bicolor_SH1509369.08FU
Mortierella_sp_SH1608153.08FU
Hyaloscypha_vraolstadiae_SH1523767.08FU
Remaining taxa (206)
% Read Abundance
50
40
30
20
10
2013
2014
2015
2016
2017
2018


(S6c)

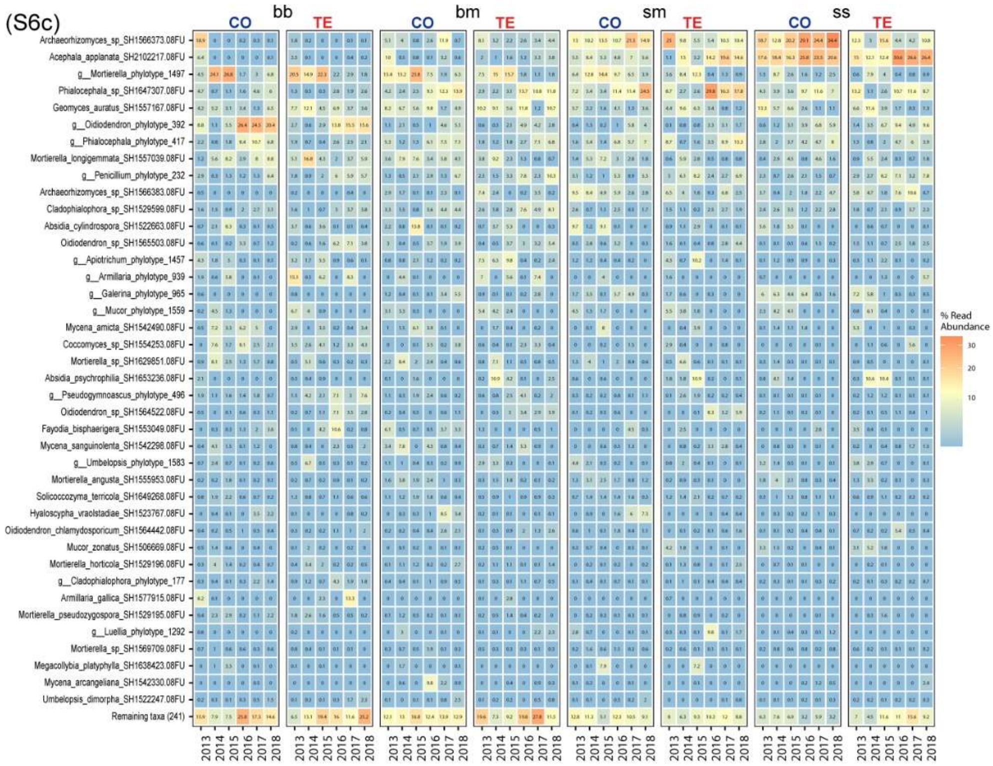

bb
bm
sm
ss
CO
TE
% Read Abundance
30
20
10
2013
2014
2015
2016
2017
2018

**Fig. S7** heatmap time-courses for fungi with unknown lifestyles in upper (S7a) and lower soil layers (S7b) and averaged for both layers (S7c); bb, bm, sm, ss: root zones (bb – pure beech, bm – beech mixture, sm – spruce mixture, ss – pure spruce); CO: control plots, TE: throughfall exclusion plots

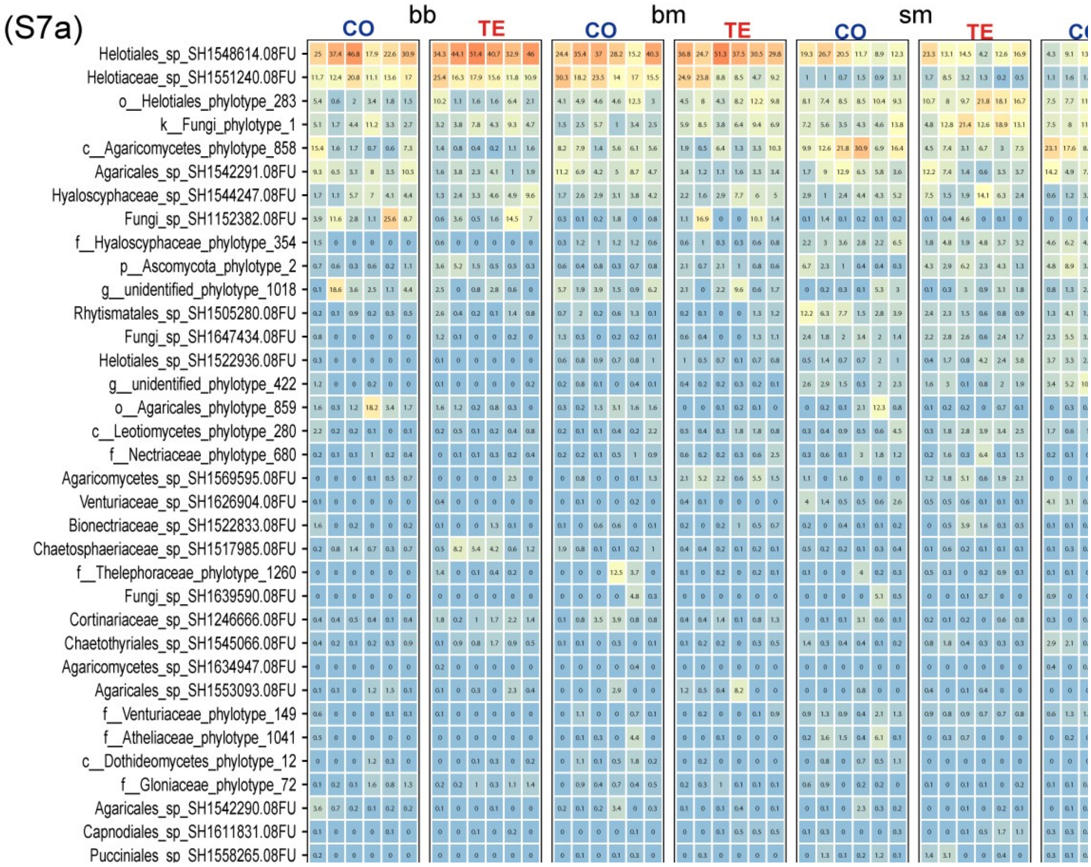

(S7b)

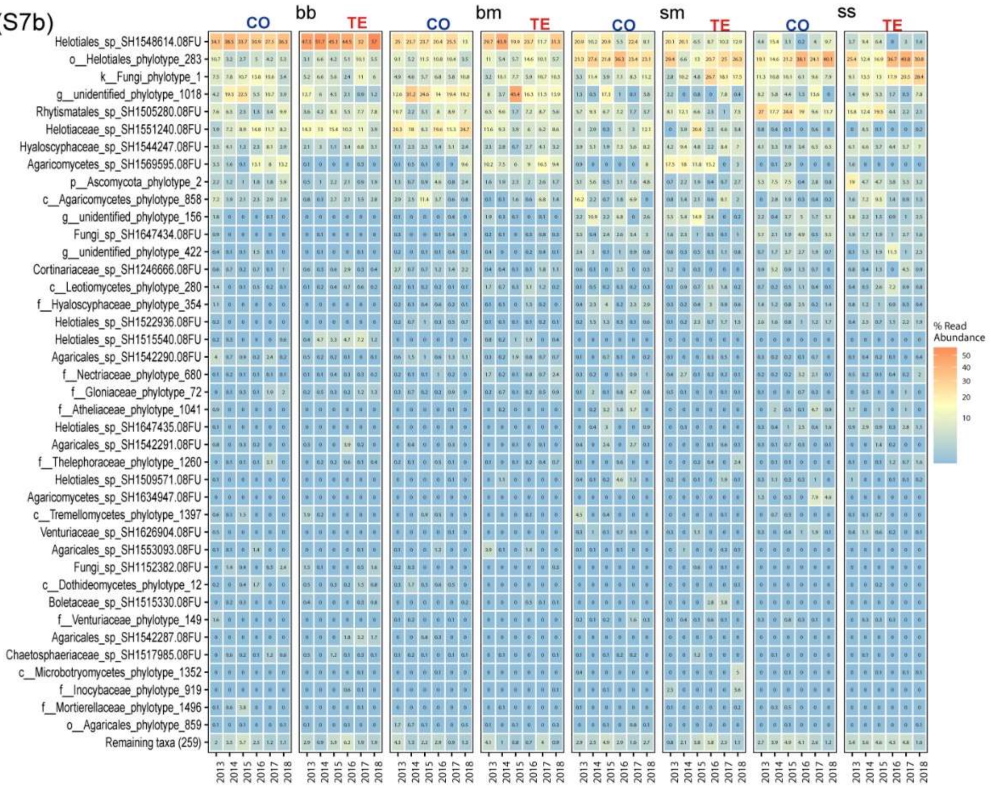
bb
bm
sm
ss
CO
TE
Helotiales_sp_SH1548614.08FU
o__Helotiales_phylotype_283
k__Fungi_phylotype_1
g__unidentified_phylotype_1018
Rhytismatales_sp_SH1505280.08FU
Helotiaceae_sp_SH1551240.08FU
Hyaloscyphaceae_sp_SH1544247.08FU
Agaricomycetes_sp_SH1569595.08FU
p__Ascomycota_phylotype_2
c__Agaricomycetes_phylotype_858
g__unidentified_phylotype_156
Fungi_sp_SH1647434.08FU
g__unidentified_phylotype_422
Cortinariaceae_sp_SH1246666.08FU
c__Leotiomycetes_phylotype_280
f__Hyaloscyphaceae_phylotype_354
Helotiales_sp_SH1522936.08FU
Helotiales_sp_SH1515540.08FU
Agaricales_sp_SH1542290.08FU
f__Nectriaceae_phylotype_680
f__Gloniaceae_phylotype_72
f__Atheliaceae_phylotype_1041
Helotiales_sp_SH1647435.08FU
Agaricales_sp_SH1542291.08FU
f__Thelephoraceae_phylotype_1260
Helotiales_sp_SH1509571.08FU
Agaricomycetes_sp_SH1634947.08FU
c__Tremellomycetes_phylotype_1397
Venturiaceae_sp_SH1626904.08FU
Agaricales_sp_SH1553093.08FU
Fungi_sp_SH1152382.08FU
c__Dothideomycetes_phylotype_12
Boletaceae_sp_SH1515330.08FU
f__Venturiaceae_phylotype_149
Agaricales_sp_SH1542287.08FU
Chaetosphaeriaceae_sp_SH1517985.08FU
c__Microbotryomycetes_phylotype_1352
f__Inocybaceae_phylotype_919
f__Mortierellaceae_phylotype_1496
o__Agaricales_phylotype_859
Remaining taxa (259)
2013
2014
2015
2016
2017
2018
% Read Abundance
50
40
30
20
10

(S7c)

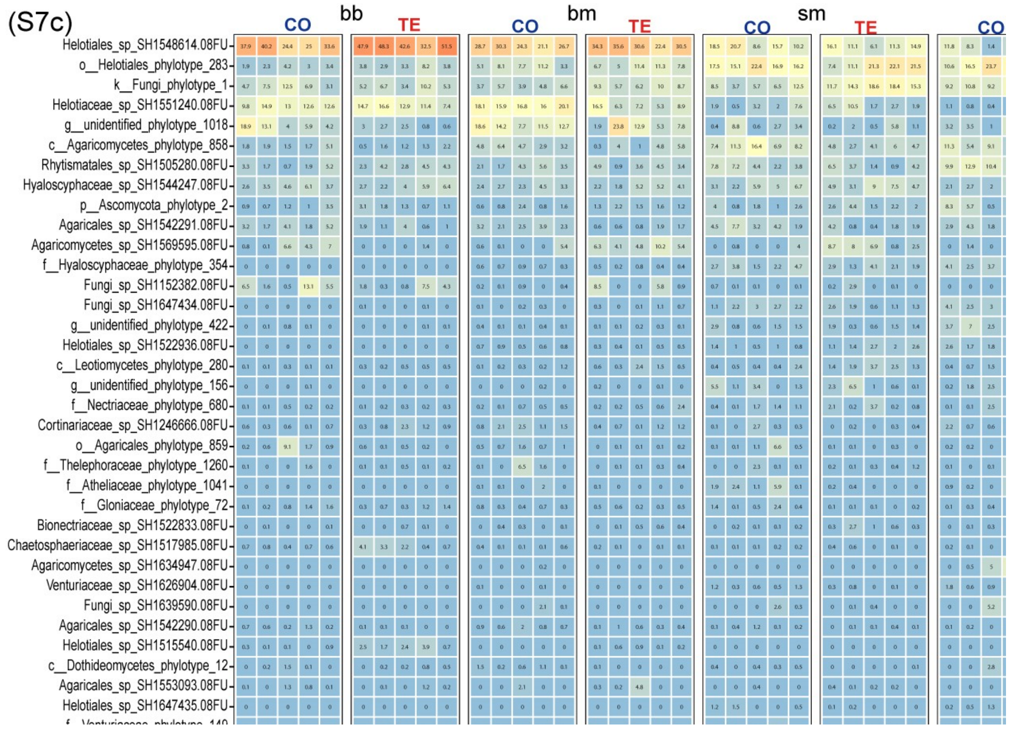
bb
bm
sm
CO
TE
Helotiales_sp_SH1548614.08FU
o__Helotiales_phylotype_283
k__Fungi_phylotype_1
Helotiaceae_sp_SH1551240.08FU
g__unidentified_phylotype_1018
c__Agaricomycetes_phylotype_858
Rhytismatales_sp_SH1505280.08FU
Hyaloscyphaceae_sp_SH1544247.08FU
p__Ascomycota_phylotype_2
Agaricales_sp_SH1542291.08FU
Agaricomycetes_sp_SH1569595.08FU
f__Hyaloscyphaceae_phylotype_354
Fungi_sp_SH1152382.08FU
Fungi_sp_SH1647434.08FU
g__unidentified_phylotype_422
Helotiales_sp_SH1522936.08FU
c__Leotiomycetes_phylotype_280
g__unidentified_phylotype_156
f__Nectriaceae_phylotype_680
Cortinariaceae_sp_SH1246666.08FU
o__Agaricales_phylotype_859
f__Thelephoraceae_phylotype_1260
f__Atheliaceae_phylotype_1041
f__Gloniaceae_phylotype_72
Bionectriaceae_sp_SH1522833.08FU
Chaetosphaeriaceae_sp_SH1517985.08FU
Agaricomycetes_sp_SH1634947.08FU
Venturiaceae_sp_SH1626904.08FU
Fungi_sp_SH1639590.08FU
Agaricales_sp_SH1542290.08FU
Helotiales_sp_SH1515540.08FU
c__Dothideomycetes_phylotype_12
Agaricales_sp_SH1553093.08FU
Helotiales_sp_SH1647435.08FU

**Table S1** Primer sequences for high throughput sequencing according to Tedersoo et al. (2015) with adapter sequences for Illumina MiSeq. References given in Materials and Methods of the main text.

| **ITS primers + adaptor sequences (MiSeq)** | |
|---|---|
| ITS3-Mix1 (Fungi) | TCGTCGGCAGCGTCAGATGTGTATAAGAGACAGCATCGATGAAGAACGCAG |
| ITS3-Mix2 (Chytridiomycota) | TCGTCGGCAGCGTCAGATGTGTATAAGAGACAGCAACGATGAAGAACGCAG |
| ITS3-Mix3 (Sebacinales) | TCGTCGGCAGCGTCAGATGTGTATAAGAGACAGCACCGATGAAGAACGCAG |
| ITS3-Mix4 (Glomeromycota) | TCGTCGGCAGCGTCAGATGTGTATAAGAGACAGCATCGATGAAGAACGTAG |
| ITS3-Mix5 (Sordariales) | TCGTCGGCAGCGTCAGATGTGTATAAGAGACAGCATCGATGAAGAACGTGG |
| ITS4-Mix1 (Fungi) | GTCTCGTGGGCTCGGAGATGTGTATAAGAGACAGTCCTCCGCTTATTGATATGC |
| ITS4-Mix2 (Chaetothyriales) | GTCTCGTGGGCTCGGAGATGTGTATAAGAGACAGTCCTGCGCTTATTGATATGC |
| ITS4-Mix3 (Archaeorhizomyc.) | GTCTCGTGGGCTCGGAGATGTGTATAAGAGACAGTCCTCGCCTTATTGATATGC |
| ITS4-Mix4 (Tulasnellaceae) | GTCTCGTGGGCTCGGAGATGTGTATAAGAGACAGTCCTCCGCTGAWTAATATGC |

**Table S2** Annual PERMANOVA results with reduced experimental units: Values from Fig. 3 (Annual PERMANOVA results on major experimental factors for important fungal groups) which used all available experimental units in comparison to values from annual models that only used four experimental units that were available during the whole course of the experiment (values in brackets). Number of experimental units was reduced during 2016 - 2018, because of dying spruce in two respective TE plots. 2013-2015: nunits=6, 2016-2018: nunits=4. Values in brackets: nunits=4 for 2013-2015). All else identical to Fig. 3 in the main text.

| **fungal group** | **factor** | **2013** pre-drought | | **2014** | | **2015** | | **2016** | | **2017** | | **2018** | |
|---|---|---|---|---|---|---|---|---|---|---|---|---|---|
| | | $R^2_{adj}$ | P | $R^2_{adj}$ | P | $R^2_{adj}$ | P | $R^2_{adj}$ | P | $R^2_{adj}$ | P | $R^2_{adj}$ | P |
| EMf (ectomycorrhizal fungi) | Treatment<br>(CO, TE) | 0.005<br>(0.007) | 0.059<br>(0.069) | **0.010**<br>(0.006) | **0.005**<br>(0.068) | **0.020**<br>**0.020** | **<.001**<br>**(0.003)** | **0.071** | **<.001** | **0.031** | **<.001** | **0.016** | **0.005** |
| | Root zone<br>(bb, bm, sm, ss) | **0.029**<br>**(0.036)** | **<.001**<br>**(0.001)** | **0.041**<br>**(0.050)** | **<.001**<br>**(<.001)** | **0.092**<br>**0.123** | **<.001**<br>**(<.001)** | **0.044** | **<.001** | **0.073** | **<.001** | **0.073** | **<.001** |
| | Soil layer of the roots<br>(upper, lower) | **0.033**<br>**(0.033)** | **<.001**<br>**(<.001)** | **0.018**<br>**(0.012)** | **<.001**<br>**(0.019)** | **0.009**<br>**0.006** | **0.009**<br>**(0.047)** | **0.009** | **0.039** | **0.015** | **0.008** | **0.025** | **<.001** |
| Saprotrophic fungi | Treatment<br>(CO, TE) | 0<br>(0) | 0.291<br>(0.43) | **0.027**<br>**(0.020)** | **<.001**<br>**(<.001)** | **0.022**<br>**(0.021)** | **<.001**<br>**(<.001)** | **0.033** | **<.001** | **0.029** | **<.001** | **0.017** | **0.002** |
| | Root zone<br>(bb, bm, sm, ss) | **0.063**<br>**(0.048)** | **<.001**<br>**(<.001)** | **0.094**<br>**(0.073)** | **<.001**<br>**(<.001)** | **0.093**<br>**(0.104)** | **<.001**<br>**(<.001)** | **0.134** | **<.001** | **0.095** | **<.001** | **0.145** | **<.001** |
| | Soil layer of the roots<br>(upper, lower) | **0.068**<br>**(0.075)** | **<.001**<br>**(<.001)** | **0.04**<br>**(0.031)** | **<.001**<br>**(<.001)** | **0.039**<br>**(0.041)** | **<.001**<br>**(<.001)** | **0.023** | **<.001** | **0.024** | **<.001** | **0.044** | **<.001** |
| Fungi of unkown lifestyle | Treatment<br>(CO, TE) | 0.004<br>(0.006) | 0.070<br>(0.065) | 0<br>(0) | 0.248<br>(0.215) | **0.006**<br>**(0)** | **0.026**<br>**(0.164)** | **0.011** | **0.016** | **0.027** | **0.002** | 0.005 | 0.057 |
| | Root zone<br>(bb, bm, sm, ss) | **0.153**<br>**(0.150)** | **<.001**<br>**(<.001)** | **0.162**<br>**(0.169)** | **<.001**<br>**(<.001)** | **0.215**<br>**(0.227)** | **<.001**<br>**(<.001)** | **0.190** | **<.001** | **0.167** | **<.001** | **0.234** | **<.001** |
| | Soil layer of the roots<br>(upper, lower) | **0.070**<br>**(0.073)** | **<.001**<br>**(<.001)** | **0.066**<br>**(0.053)** | **<.001**<br>**(<.001)** | **0.050**<br>**(0.045)** | **<.001**<br>**(<.001)** | **0.048** | **<.001** | **0.036** | **<.001** | **0.056** | **<.001** |

**Table S3** PERMANOVA results for Fig. 4b,d,f (results of the function adonis() (vegan 2.5.7) in models including all significant interactions, omitted in the Fig. 4). All else identical to Fig. 4b,d,f.

| **EMf** | Df | SumsOfSqs | MeanSqs | F.Model | R2 | Pr(>F) |
|---|---|---|---|---|---|---|
| Treatment (CO, TE) | 1 | 3.5672 | 3.5672 | 17.4288 | 0.0450 | 0.0001 |
| Root zone (bb, bm, sm, ss) | 3 | 8.0673 | 2.6891 | 13.1384 | 0.1017 | 0.0001 |
| Soil layer of the roots (upper, lower) | 1 | 1.9553 | 1.9553 | 9.5535 | 0.0246 | 0.0001 |
| Year of treatment (2014-2018) | 1 | 1.0105 | 1.0105 | 4.9371 | 0.0127 | 0.0001 |
| Vital root tips per cm³ soil | 1 | 0.6463 | 0.6463 | 3.1576 | 0.0081 | 0.0859 |
| $N_{min}$ in the sampled soil | 1 | 0.4807 | 0.4807 | 2.3487 | 0.0061 | 0.0115 |
| Treatment × Root zone | 3 | 1.3053 | 0.4351 | 2.1258 | 0.0164 | 0.0002 |
| Treatment × Soil layer of the roots | 1 | 0.6255 | 0.6255 | 3.0561 | 0.0079 | 0.0009 |
| Root zone × Vital root tips | 3 | 0.7234 | 0.2411 | 1.1782 | 0.0091 | 0.0395 |
| Soil layer of the roots × Vital root tips | 1 | 0.5049 | 0.5049 | 2.4667 | 0.0064 | 0.0157 |
| Treatment × Nmin | 1 | 0.9090 | 0.9090 | 4.4413 | 0.0115 | 0.0047 |
| *Residuals* | *291* | *59.5599* | *0.2047* | | *0.7505* | |
| *Total* | *308* | *79.3553* | | | *1.0000* | |

| **Saprotroph** | Df | SumsOfSqs | MeanSqs | F.Model | R2 | Pr(>F) |
|---|---|---|---|---|---|---|
| Treatment (CO, TE) | 1 | 1.9167 | 1.9167 | 13.2604 | 0.0300 | 0.0001 |
| Root zone (bb, bm, sm, ss) | 3 | 7.1420 | 2.3807 | 16.4700 | 0.1118 | 0.0001 |
| Soil layer of the roots (upper, lower) | 1 | 4.0027 | 4.0027 | 27.6916 | 0.0627 | 0.0001 |
| Year of treatment (2014-2018) | 1 | 5.0671 | 5.0671 | 35.0554 | 0.0793 | 0.0001 |
| Vital root tips per cm³ soil | 1 | 0.6680 | 0.6680 | 4.6211 | 0.0105 | 0.0001 |
| $N_{min}$ in the sampled soil | 1 | 0.1864 | 0.1864 | 1.2897 | 0.0029 | 0.1816 |
| Root zone × Soil layer of the roots | 3 | 0.7497 | 0.2499 | 1.7288 | 0.0117 | 0.0016 |
| Root zone × Year of treatment | 3 | 0.6098 | 0.2033 | 1.4061 | 0.0095 | 0.0257 |
| Treatment × Year of treatment | 1 | 0.4712 | 0.4712 | 3.2599 | 0.0074 | 0.0003 |
| Soil layer × Year of treatment | 1 | 0.3246 | 0.3246 | 2.2460 | 0.0051 | 0.0043 |
| Soil layer × Vital root tips | 1 | 0.2637 | 0.2637 | 1.8242 | 0.0041 | 0.0358 |
| Year of treatment × Vital root tips | 1 | 0.2904 | 0.2904 | 2.0091 | 0.0045 | 0.0119 |
| Treatment × Nmin | 1 | 0.4194 | 0.4194 | 2.9016 | 0.0066 | 0.0014 |
| *Residuals* | *289* | *41.7734* | *0.1445* | | *0.6539* | |
| *Total* | *308* | *63.8851* | | | *1.0000* | |

| **unknown lifestlye** | Df | SumsOfSqs | MeanSqs | F.Model | R2 | Pr(>F) |
|---|---|---|---|---|---|---|
| Treatment (CO, TE) | 1 | 0.8310 | 0.8310 | 7.4639 | 0.0161 | 0.0001 |
| Root zone (bb, bm, sm, ss) | 3 | 10.0300 | 3.3433 | 30.0306 | 0.1948 | 0.0001 |
| Soil layer of the roots (upper, lower) | 1 | 3.8274 | 3.8274 | 34.3788 | 0.0743 | 0.0001 |
| Year of treatment (2014-2018) | 1 | 0.6175 | 0.6175 | 5.5465 | 0.0120 | 0.0001 |
| Vital root tips per cm³ soil | 1 | 0.7497 | 0.7497 | 6.7344 | 0.0146 | 0.0001 |
| $N_{min}$ in the sampled soil | 1 | 0.1930 | 0.1930 | 1.7337 | 0.0037 | 0.0473 |
| Treatment × Root zone | 3 | 0.8176 | 0.2725 | 2.4478 | 0.0159 | 0.0001 |
| Root zone × Soil layer of the roots | 3 | 1.2285 | 0.4095 | 3.6782 | 0.0239 | 0.0001 |
| Treatment × Soil layer of the roots | 1 | 0.2748 | 0.2748 | 2.4683 | 0.0053 | 0.0056 |
| Treatment × Year of treatment | 1 | 0.2659 | 0.2659 | 2.3883 | 0.0052 | 0.0070 |
| Root zone × Nmin | 3 | 0.4775 | 0.1592 | 1.4296 | 0.0093 | 0.0415 |
| *Residuals* | *289* | *32.1746* | *0.1113* | | *0.6249* | |
| *Total* | *308* | *51.4874* | | | *1.0000* | |

**Methods S1** Ectomycorrhizal enzyme activity tests

Potential extracellular enzyme activities (EAs) were determined on individual root tips sequentially incubated with eight enzyme substrates as described by Pritsch et al. (2011). Vital tips assigned to morphotypes were assayed, and enzyme activities were calculated per tip ($EA_{tip}$) and per volume of the soil sample ($EA_{vol}$) as described by Nickel et al. (2018). In brief, 21 fully vital ECM tips were assayed representing the share of ECM morphotypes (Agerer 2001) (n ≥ 3) in each sample based on vital tip counting. We used seven fluorescent enzyme substrates based on 4-methylumbelliferone (MU), respectively 7-aminomethylcoumarin (AMC), and 2,2'-azino-bis(3-ethylbenzothiazoline-6-sulphonic acid) (ABTS) in a 96-well microplate assay. The following enzyme activity potentials were tested: leucine aminopeptidase (EC 3.4.11.1), xylosidase (EC 3.2.1.37), glucuronidase (EC 3.2.1.31), cellobiohydrolase (EC 3.2.1.91), N-acetyl-glucosaminidase (EC 3.2.1.14), β-glucosidase (EC 3.2.1.3), phosphatase (EC 3.1.3.2) and laccase (EC 1.10.3.2). These data were used to calculate the average EA per tip of each sample ($EA_{tip}$) as the weighted mean of assayed tips. Based on the total number of vital tips per soil volume, we also calculated $EA_{vol}$ taking into account the recorded soil volume of each sample.

Agerer, R. (2001). Exploration types of ectomycorrhizae. *Mycorrhiza, 11*(2), 107-114.

Nickel, U. T., Weikl, F., Kerner, R., Schafer, C., Kallenbach, C., Munch, J. C., & Pritsch, K. (2018). Quantitative losses vs. qualitative stability of ectomycorrhizal community responses to 3 years of experimental summer drought in a beech-spruce forest. *Glob Chang Biol, 24*(2), e560-e576. doi:10.1111/gcb.13957

Pritsch, K., Courty, P. E., Churin, J. L., Cloutier-Hurteau, B., Ali, M. A., Damon, C., . . . Garbaye, J. (2011). Optimized assay and storage conditions for enzyme activity profiling of ectomycorrhizae. *Mycorrhiza, 21*(7), 589-600. doi:10.1007/s00572-011-0364-4
